# *Leishmania donovani* GP63 paralogues cooperatively orchestrate visceral infection and persistence

**DOI:** 10.1101/2025.11.09.687440

**Authors:** Debolina Manna, Sunandini Bhattacharya, Shounak Jana, Kristína Záhonová, Suvendu Nandi, Deneshraj Srinivasan, Nilanjan Pradhan, Nidhi Pandey, Debarati Biswas, Mahitosh Mandal, Gayatri Mukherjee, Vyacheslav Yurchenko, Budhaditya Mukherjee

**Affiliations:** School of Medical Science and Technology, Indian Institute of Technology Kharagpur, Kharagpur, India; Life Science Research Centre, Faculty of Science, University of Ostrava, Ostrava, Czech Republic; Institute of Parasitology, Biology Centre, Czech Academy of Sciences, České Budějovice, Czech Republic; Department of Parasitology, Faculty of Science, Charles University, BIOCEV, Vestec, Czech Republic; Division of Infectious Diseases, Department of Medicine, University of Alberta, Edmonton, Canada

**Keywords:** Visceral leishmaniasis, metalloprotease GP63 paralogues, adaptive virulence, evolutionary divergence, host invasion and persistence, pyroptosis

## Abstract

Highly plastic genome of *Leishmania* is thought to underlie its remarkable ability to adapt to diverse host microenvironments, yet how this plasticity translates into dermal versus visceral persistence remains unclear. GP63, a zinc metalloprotease, is a well-established determinant of promastigote infectivity, particularly in cutaneous leishmaniasis. Despite this, its role in amastigote stage and in visceral disease caused by *Leishmania donovani* remains undefined. Here, using an integrated approach combining phylogenetics, CRISPR-mediated genome editing, and biochemical analyses, we demonstrate that GP63 paralogues in *L. donovani* have undergone independent evolutionary diversification to support visceral infection. While GP63-paralogues encoded on chromosome 10, previously implicated in cutaneous manifestation caused by *L. major*, were found severely truncated and dispensable in *L. donovani,* two distinct GP63-paralogues encoded on chromosome 28 and 31 were found ‘essential’ to establish visceral infection. LdGP63_31 facilitates promastigote attachment to macrophages by promoting lipid raft engagement and complement inactivation, thereby enabling host entry and subsequent amastigote differentiation. LdGP63_28 is essential for intracellular amastigote survival by suppressing host inflammatory pyroptosis. Structural and enzymatic analyses revealed preferential host localization and substrate specificities, possibly resulting from distinct amino acid substitution within conserved motifs of these proteases, indicating independent evolutionary adaptations. Importantly, loss of the LdGP63_28 more severely impaired amastigote reinfection than loss of its counterpart LdGP63_31. Together, these findings reveal coordinated functional specialization of GP63-paralogues ensuring visceral adaptation.

## INTRODUCTION

Leishmaniasis, a neglected vector-borne parasitic disease caused by the kinetoplastid protists of the genus *Leishmania* and transmitted by sand flies, endangers over a billion people across endemic regions. Its clinical manifestations range from the benign cutaneous lesions in the cases of cutaneous leishmaniasis (CL) to often fatal visceral leishmaniasis (VL) with prevalence up to 90,000 cases yearly [1, 2].

*Leishmania* parasites navigate through a dixenous life cycle shuttling between flagellated promastigotes thriving in the sand fly midgut and amastigotes replicating intracellularly within mammalian macrophages (M□) [3, 4]. So far, only a few parasite-specific virulence factors, namely lipophosphoglycan (LPG), glycoinositol phospholipid (GIPL), and a zinc metalloprotease GP63 (glycoprotease 63 or leishmanolysin), have been implicated in enabling parasites’ attachment, invasion, complement evasion, and survival in the host [5–7]. During initial infection in mammalian host, GP63 expressed by *Leishmania* promastigotes helps parasites to evade lysis by inactivating the host complement component C3b opsonizing the entering parasites and allowing them to bind to the host-macrophage (M□) membrane receptors CR1/CR3 [8–10]. Following attachment, invading parasites extract cholesterol from M□ lipid rafts, a process that facilitates efficient membrane remodelling and completion of host cell entry [11]. Once internalized, promastigotes differentiate into amastigotes within parasitophorous vacuoles (PVs) [12], where intracellular survival is supported by multiple parasite-derived mechanisms, including the secretion of virulence-associated proteases such as GP63 [13]. Later, these proteases hijack host signalling pathways to facilitate lysis of infected M□ enabling parasite dissemination and secondary infection [14]. While the role of GP63 in modulating *Leishmania* entry and hijacking of host pathways is well studied [15], its importance, if any, in governing amastigote persistence and dissemination within the mammalian host remain poorly defined.

Notably, the *gp63* gene family exemplifies evolutionary divergence, forming one of *Leishmania*’s largest and most polymorphic gene clusters across chromosomes 10, 28, and 31 (two in chromosome 10, one in chromosome 28, and one in chromosome 31 for *Leishmania donovani*) [16, 17]. The GP63 paralogues vary significantly among different *Leishmania* spp. in terms of copy number, which varies from none to as high as 33 within chromosomes of some species [18–20]. This remarkable variability suggests lineage-specific adaptive divergence driven not only by the host species, but also by the distinct host microenvironments, in which the parasites persist [21–23]. The studies in CL-causing *L. major* and *L. amazonensis* have demonstrated that parasites lacking *gp63* gene copies encoded on chromosome 10 induce delayed lesion development in infected mice compared with wild-type infection, yet, these mutants retain substantial residual virulence [24, 25]. However, the role of GP63 in VL-causing *Leishmania* spp. remains unexplored, with some reports suggesting a drastic drop in GP63 expression in amastigotes of some species [26, 27].

Here, we document that GP63 paralogues function cooperatively and distinctly to drive infectivity and host persistence in visceral microenvironment. Two paralogues on chromosome 10 that were previously implicated in CL proved redundant for visceral infection. Conversely, a paralogue encoded on chromosome 28 was found indispensable for amastigote survival, acting by suppressing inflammasome activation and inhibiting host pyroptosis. In contrast, a paralogue encoded on chromosome 31 was documented to govern *L. donovani* attachment to lipid rafts and inactivation of host C3b ensuring successful parasite entry. This work thus addresses how GP63 diversification, through independent evolution, might have enabled stage-specific functioning in the visceral context, eventually giving rise to independent GP63 paralogues having different chromosomal origins that play prominent roles in *L. donovani* infection.

## RESULTS

### 1. *L. donovani* possesses four GP63 paralogues

In *L. donovani*, four GP63 paralogues were found in the TriTrypDB [28], two encoded on chromosome (chr-) 10 (*LdBPK_100510.1* [*LdGP63_10.51*] and *LdBPK_100520.1* [*LdGP63_10.52*]), one on chr-28 (*LdBPK_280600.1* [*LdGP63_28*]), and one on chr-31 (*LdBPK_312040.1* [*LdGP63_31*]). No additional paralogues were identified by BLAST searches. Both paralogues on chr-10 appeared to be significantly truncated (only 85 and 86 amino acids long), whereas the other two paralogues were of sizes similar to those documented in CL causing *Leishmania major* (566 and 636 amino acids) (**Fig. 1A**). Importantly, phylogenetic inference, using full-length GP63 sequences and including GP63 from selected Leishmaniinae spp. and *Angomonas deanei* as an outgroup (**Table S1**), resolved three highly-supported clades comprising GP63 encoded on different chromosomes (**Fig. 1B, Fig. S1**), namely chr-10, 28, and 31. *L. donovani* GP63 of chr-28 and 31 clustered with another VL-causing strain, *Leishmania infantum*, further highlighting that these GP63 likely remain evolutionary conserved.

**Figure 1:**
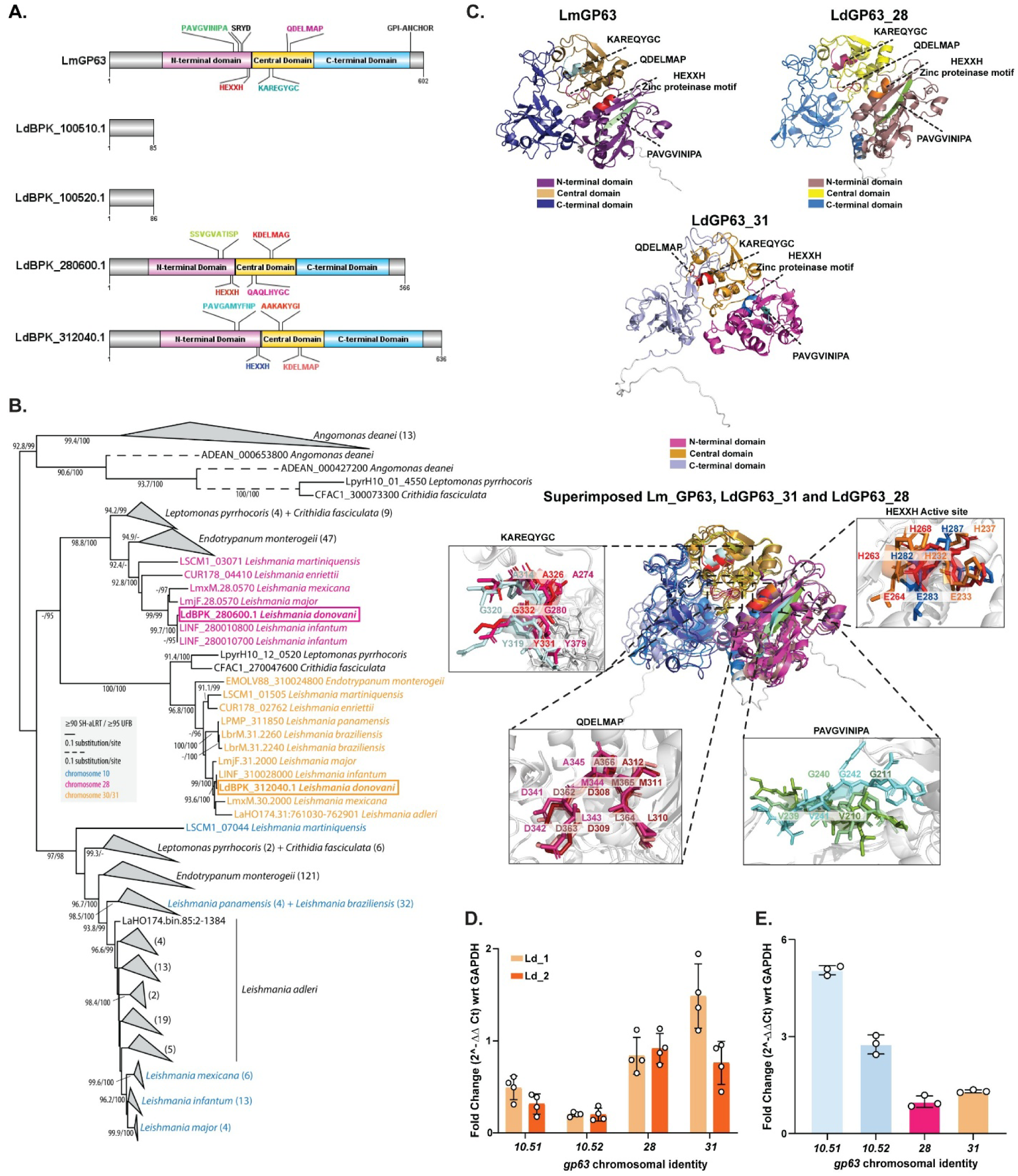
*LdGP63_28* and *LdGP63_31* represent functionally active GP63 proteases in *L. donovani*. A. Structural comparisons among one *L. major* (Lm) GP63 and *L. donovani* (Ld) GP63 sequences. B. Phylogenetic analysis of selected Leishmaniinae GP63. *L.donovani LdGP63_28* and *LdGP63_31* are enclosed in pink and orange boxes, respectively. Some clades were collapsed for visualization purposes; for the full tree see **Fig. S1**. C. Crystal structure of LmGP63 (PDB: 1LML) and AlphaFold models of LdGP63_28 and LdGP63_31, followed by superimposed structure of all three proteases showing overlaps of the HEXXH motif and conserved regions (PAVGVINIPA, KAREQYGC, and QDELMAP), within the N-terminal (LmGP63: magenta, LdGP63_28: brown, LdGP63_31: pink), central (LmGP63: ochre, LdGP63_28: light yellow, LdGP63_31: golden yellow), and C-terminal (LmGP63: dark blue, LdGP63_28: royal blue, LdGP63_31: lavender) domains (the insets show detailed view of the overlapping conserved regions with their common amino acid residues). D, E. qPCR analysis showing expression of *LdGP63_28* and *LdGP63_31* in two independent *L. donovani* strains (Ld_1 and Ld_2) in promastigotes (D) and intracellular amastigotes (E) in infected murine M□ at 24h p.i. Data are presented as mean ± standard error, with each individual point (n = 4 for D and n = 3 for E) shown. Statistical analysis was performed by paired two-tailed student’s *t*-test: ns, non-significant (P > 0.05); *, P ≤ 0.05; **, P ≤ 0.01; ***, P ≤ 0.001; ****, P ≤ 0.0001.

Given that the chr-10-encoded GP63 from *L. major* (LmGP63; LmjF.10.0460) has been extensively characterized [29], we compared its protein sequence and structural features with GP63 paralogues from *L. donovani*. Multiple sequence alignment revealed strong conservation of the N-terminal, central, and C-terminal domains, as well as the conserved sequence motifs (PAVGVINIPA, KAREQYGC, QDELMAP) and the catalytic core HEXXH [17, 29], in both LdGP63_28 and LdGP63_31, which were expectedly absent in the truncated LdGP63_10.51 and LdGP63_10.52 (**Fig.1A, Fig. S2A**).

To assess structural conservation, AlphaFold-predicted models of LdGP63_28 and LdGP63_31 were superimposed onto the crystal structure of LmGP63 (PDB: 1LML) (**Fig. 1C**). Both paralogues exhibited favourable Ramachandran plot distributions (**Fig. S2B**), with substantial structural similarities having TM-scores of 0.77998 and 0.645 19 (>0.5) and RMSD values of 1.763 Å and 3.499 Å (<4 Å), respectively, indicating a conserved overall fold and substantial overlap of key structural residues. Interestingly, we also observed distinct amino acid substitutions among these conserved motifs of LdGP63_28, LdGP63_31 and LmGP63 (**Fig. 1C**).

Consistent with these observations, RT–qPCR analysis confirmed robust expression of both *LdGP63_28* and *LdGP63_31* in promastigote stages across two independent clinical *L. donovani* isolates (**Fig. 1D**), as well as in intracellular amastigotes (**Fig. 1E**). However, surprisingly, expression of *LdGP63_10.51* and *LdGP63_10.52* was also observed in both the cases (**Fig. 1D, E**).

### 2. LdGP63_10.51 and LdGP63_10.52 are dispensable, non-functional proteases

Since *LdGP63* from chr-10 showed detectable expression at RNA level (**Fig. 1D, E**), we wanted to further investigate the chr-10 paralogues despite their fragmented nature. For functional characterization, Cas9-expressing *L. donovani* lines were generated and confirmed (**Fig. S3A, B**), and deletion of *LdGP63_10.51* and *LdGP63_10.52* were performed separately in these lines. Both *LdGP63_10.51* and *LdGP63_10.52* knockouts (KOs) (**Fig. S3C, D**) revealed no defects in growth, morphology, or motility relative to the wild-type (WT) or parental lines (**Fig. S3E, F**), nor resulted in any impairment in metacyclogenesis (**Fig. S3G-J**). Moreover, comparative attachment assays [30] in host M□ demonstrated that both the LdGP63_10.51_KO and LdGP63_10.52_KO lines attach to host M□ as efficiently as parental (**Fig. S3K, L**), resulting in equivalent intracellular amastigote loads at 6□h, 24□h, and 48□h□post infection (p.i.) (**Fig. S3M, N**). The truncated *LdGP63_10.51* and *LdGP63_10.52* devoid of catalytic motifs were thus confirmed to be dispensable for promastigotes and M□ infection.

### 3. LdGP63_28 is essential for amastigote persistence in *L. donovani* infected M□

To investigate possible function of *LdGP63_28*, previously generated Cas9-expressing *L. donovani* lines (**Fig. S3A, B**) were used to perform C-terminal endogenous tagging of *LdGP63_28* with mCherry (**Fig.□2A, B**). In parallel, LdGP63_28_KO lines were also generated (**Fig. S4A**) in the same Cas9 background and confirmed by PCR (**Fig. S4B**). Ultrastructure-expansion of LdGP63_28mCherry promastigotes, revealed LdGP63_28 to have a punctuated vesicular distribution preferentially localized near the flagellar pocket (**Fig. 2C**), indicative of its possible vesicular secretion. Consistent with this, confocal microscopy showed that LdGP63_28 co-localized with Rab5a positive secretory vesicles in promastigotes (**Fig. 2D; Video S1**). During M□ infection, LdGP63_28 was detected within host endosomal and exosomal compartments (**Fig. 2E; Video S2**), localizing with Rab5a, CD63, and CD9 positive vesicles. Notably, LdGP63_28-containing vesicles were observed both within infected M□ and among extracellular vesicles released into the surrounding environment, confirming its secretory nature (**Fig. 2E**), similar to what have been reported for LmGP63 [31]. Furthermore, LdGP63_28 was also predicted to harbour a signal peptide (Probability = 0.823815), with comparable prediction score (Probability = 0.671865) as LmGP63 (**Table S2**). Although, LdGP63_28_KO lines also displayed no defect in growth, morphology, motility (**Fig. S4C, D; Video**□**S3**), or metacyclogenesis (**Fig. S4E, F**), LdGP63_28_KO promastigotes showed a modestly, yet statistically significant, reduced host attachment, compared to the parental line (**Fig. 2F, G**). Surprisingly, more pronounced defect was observed in intracellular amastigotes proliferation with LdGP63_28_KO amastigotes number declining sharply by 24□h p.i. and almost disappearing by 48□h□p.i. (**Fig. 2H, I**). This defect in LdGP63_28_KO amastigote proliferation was also accompanied by a decrease in the number of host M□ (**Video**□**S4**), with the existing ones showing a granular and shrunken phenotype (**Fig. S4G, H**). Annexin-V/Propidium iodide staining, which determines dead vs live M□ population, revealed increased population of viable M□ in parental infection with respect to LdGP63_28_KO infection (**Fig.**□**S4I**), indicating a probable inflammatory death of host M□ in the absence of LdGP63_28. Complementation with parental full-length copy of LdGP63_28-eGFP (LdGP63_28 addback) restored host attachment (**Fig. 2F**, right panel), and vesicular cytosolic localization of LdGP63_28-eGFP in host M□ (**Fig. 2H**, 3^rd^ and 4^th^ panels), thereby, rescuing amastigote persistence (**Fig. 2I**). Altogether, LdGP63_28 proved “essential” for *in vitro* promastigote adhesion and survival of intracellular amastigotes.

**Figure 2:**
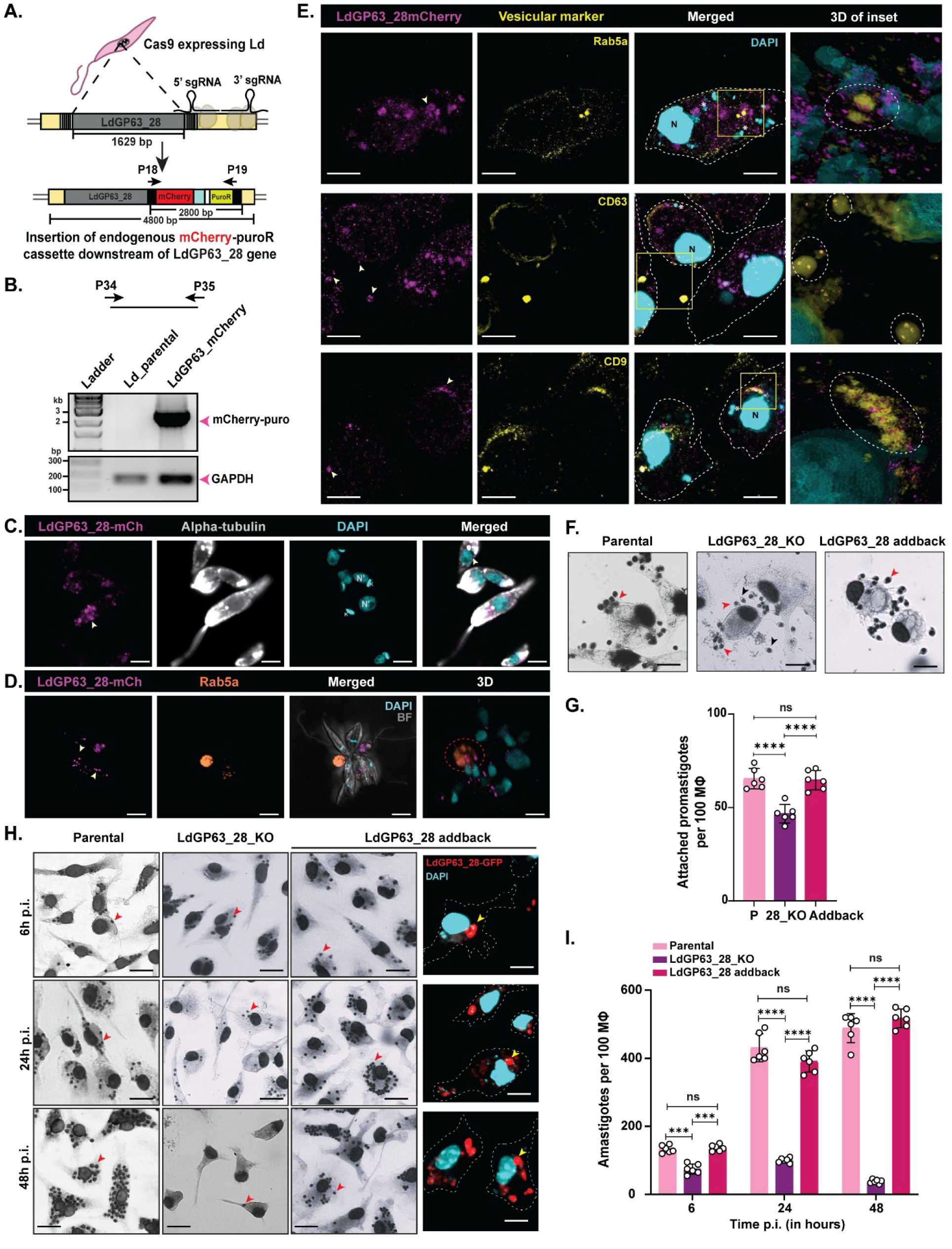
LdGP63_28 plays an essential role in infectivity and intracellular persistence of *L. donovani* amastigotes. A. Scheme depicting the endogenous tagging of mCherry-puromycin (PuroR) cassette to the C-terminal end of LdGP63_28. Primers for amplification (P18 and P19) are listed in **Table S4**. B. PCR confirmation of inserted mCherry cassette downstream of LdGP63_28. GAPDH was used as the housekeeping control. Primers for KO confirmation (P34 and P35) are listed in **Table S4**. C. Ultrastructure expansion followed by confocal imaging of promastigotes to visualize LdGP6_28mCherry (white arrows; LdGP63_28mCh) with respect to alpha-tubulin in promastigotes. DAPI (cyan) stained nucleus is denoted as N’ and kinetoplast is denoted as K. Scale bar: 2 µm. D. Representative confocal micrograph showing localization of LdGP63_28mCherry (white arrows, left panel) with Rab5a in promastigotes. The extreme right panel indicates 3D view (generated in LAS X software) of the merged image. Dotted circles represent LdGP63_28mCherry localizing with Rab5a. Scale bar: 5 µm. E. Representative confocal micrograph showing localization of LdGP63_28mCherry (white arrows in left panel) with M□ vesicular markers Rab5a, CD63, and CD9 in infected M□ at 6h p.i. White dotted line represents M□ membrane. Asterisk (*) corresponds to one amastigote nucleus (small cyan dots) and nucleus of M□ is indicatfed by N. The extreme right panel indicates 3D view (generated in LAS X software) of the merged inset region, and co-localized vesicular marker and LdGP63_28mCherry is enclosed in dotted ellipses. Scale bar: 5 µm. F. Giemsa-stained images comparing initial attachment at 30 min p.i. Attached and free *L. donovani* promastigotes are pointed by red and black arrows, respectively. Scale bar: 5 µm. G. Bar graph representing number of attached promastigotes in parental (P), LdGP63_28_KO (28_KO), and LdGP63_28 addback (Addback) lines (n = 6). H. Giemsa-stained images of murine M□ infected with parental, LdGP63_28_KO, and LdGP63_28 addback amastigotes at 6 h, 24 h, and 48 h p.i. Intracellular amastigotes and LdGP63_28-eGFP vesicles (extreme right panel) are pointed by red and yellow arrows, respectively. Scale bar: 10 µm. I. Bar graph showing the infection and proliferation of parental, LdGP63_28_KO, and LdGP63_28 addback amastigotes (n = 6). Data are presented as mean ± standard error, with each individual point shown. Statistical analysis was performed by one-way ANOVA (G) followed by Tukey’s MCT and two-way ANOVA followed by Sidak’s MCT (I): ns, non-significant (P > 0.05); *, P ≤ 0.05; **, P ≤ 0.01; ***, P ≤ 0.001; ****, P ≤ 0.0001.

### 4. LdGP63_28 deficiency triggers pyroptosis in infected host

Involvement of pyroptosis in LdGP63_28_KO infected M□ was further investigated by structured illumination super resolution microscopy (SIM^2^) [32] to precisely monitor cellular and subcellular membrane architecture. Although, at 6 h p.i., membranes and nuclei remained intact in both parental and KO infected M□, by 48 h p.i. progressive membrane fragmentation and occasional nuclear rupture were evident in response to KO infection (**Fig.**□**3A**). This punctate membrane disintegration implicated pyroptosis over apoptotic death of the infected M□. Although the number of LdGP63_28_KO infected M□ at 48□h□p.i. was low for flow analysis, at 24 h p.i., LdGP63_28_KO infected M□ revealed elevated Annexin V populations in comparison with uninfected and parental controls (**Fig. S5A, B**), suggesting concurrent apoptosis along with pyroptosis. Moreover, qPCR of LdGP63_28_KO infected M□ at 48 h p.i. confirmed marked upregulation of *nlrp3*, *caspase*lJ*3*, *caspase*lJ*2*, *caspase*lJ*7*, *caspase*lJ*1*, and to some extent *aim2*, but not *nlrc4* (**Fig.**□**S5C-J**), consistent with signs of both apoptotic and pyroptotic host death. Notably, *nlrp1* was upregulated in both infection types, but declined over time in parental infections while persisting in KO infections, suggesting a potential role of *nlrp1* to counter initial stages of *L. donovani* infection. Western blots confirmed NLRP3 elevation and cleavage of Caspase-1/3 plus Gasdermin E at 24 and 48 h p.i. in KOs (**Fig. 3B**), favouring pyroptosis alongside some possible apoptosis. Further, presence of cleaved IL-1β even at 48 h p.i. in KO-infected M□, which get supressed in parental infection, further supported NLRP3-driven sustained pyroptosis over apoptosis. Finally, SEM imaging showed membrane disintegration and “pyroptotic bodies” without significant nuclear fragmentation (**Fig. 3C, D)**, confirming host M□ mostly suffer pyroptotic death in the absence of LdGP63_28.

**Figure 3:**
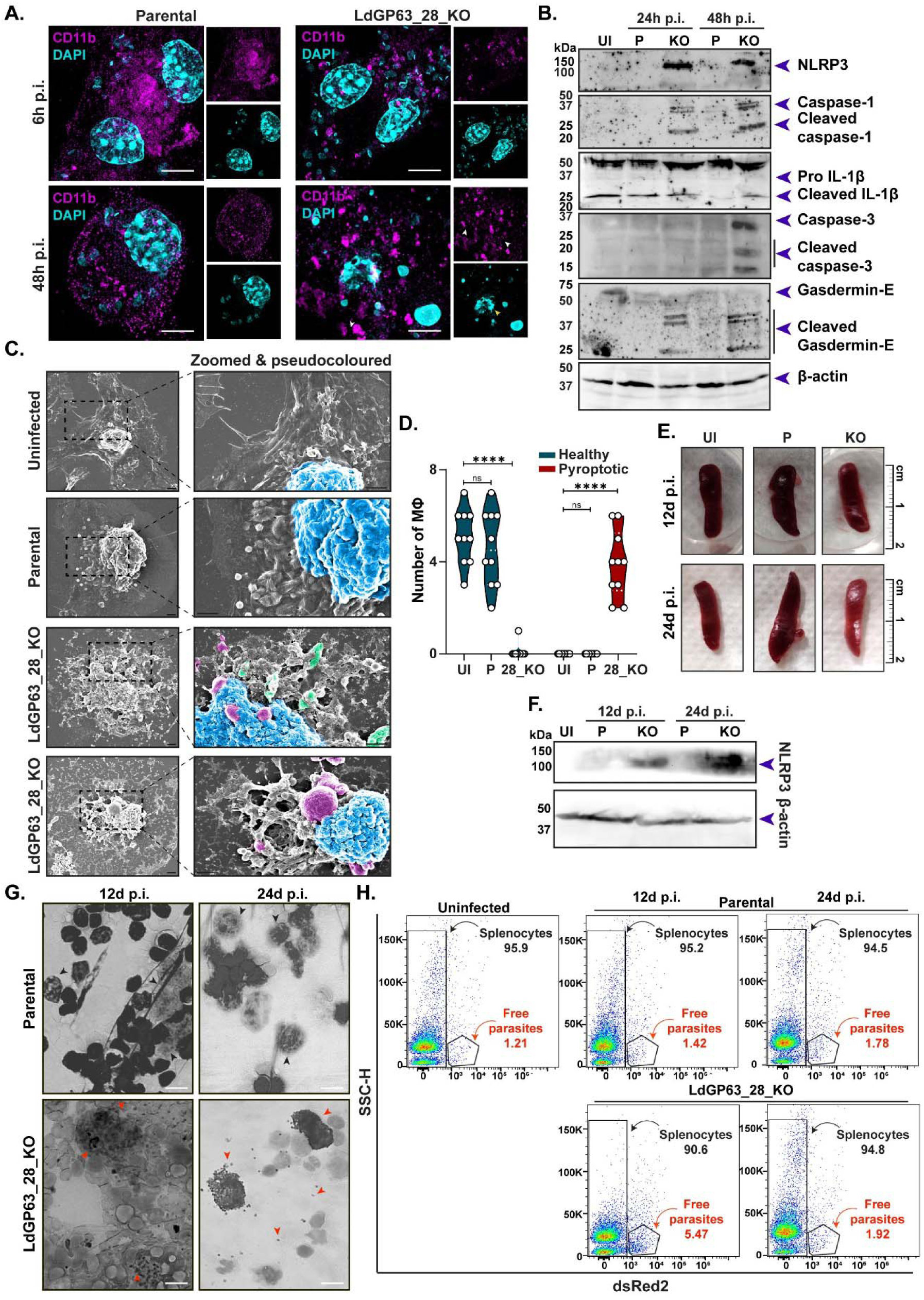
LdGP63_28 regulates pyroptosis by dampening host inflammasomal responses. A. Representative structural illumination microscopy (SIM^2^) images showing the presence of CD11b in parental and LdGP63_28_KO infected M□ at 6 h and 48 h p.i. White and yellow arrows represent fragmented M□ plasma membrane and occasional fragmented nuclear membrane, respectively. Scale bar: 2 µm. B. Western blot quantifying pyroptotic markers: NLRP3, caspase 1, IL-1β, caspase 3, and Gasdermin-E in parental (P) and LdGP63_28_KO (KO) infected M□. β-actin was used as the housekeeping control. C. SEM images of 48 h p.i. uninfected, parental and KO infected M□ showing nuclear region (blue), pyroptotic bodies (violet), and dead parasites (green). Scale bar: 2 µm. D. Graph showing number of healthy and pyroptotic M□ counted across 10 SEM images (n=10) each for uninfected (UI), parental (P), and LdGP63_28_KO (28_KO) lines. E. Spleens from uninfected (UI), parental (P), and LdGP63_28_KO (KO) infected Balb/C mice isolated at 12 days and 24 days p.i. Animal experiments involved 3 mice in each group. F. Western blot of the isolated splenocytes showing upregulation of NLRP3 in LdGP63_28_KO infected Balb/C mice. β-actin was used as the housekeeping control. G. Giemsa-stained stamp-smears of the infected spleens. Black and red arrows mark parental and LdGP63_28_KO amastigotes, respectively. Scale bar: 10 µm. H. Flow cytometric plot of Side Scatter Height (SSC-H) vs dsRed2 showing free parasites (shown by red arrow) in isolated splenocytes. Data are presented as mean ± standard error, with each individual point shown. Statistical analysis was performed by two-way ANOVA followed by Tukey’s MCT (D): ns, non-significant (P > 0.05); *, P ≤ 0.05; **, P ≤ 0.01; ***, P ≤ 0.001; ****, P ≤ 0.0001.

Primary role of LdGP63_28 in persistence of visceral amastigotes rather than parasite entry was further substantiated in murine model of VL. While LdGP63_28_KO infections in BALB/c mice did not promote significant splenomegaly (**Fig. 3E**), it significantly induced splenic expression of NLRP3 when compared to infection with parental line at 12 and 24 days p.i. (**Fig. 3F**), suggesting a possible initial infection followed by enhanced inflammatory clearance. Surprisingly, flow cytometric analysis revealed increased dsRed2⁺ splenic parasite signals at 12 days p.i. in LdGP63_28_KO-infected mice. However, by 24 days p.i., parasite burden was reduced to approximately half, whereas the parental strain showed a marked increase (**Fig. S5K**). This apparent early accumulation likely reflects pyroptotic lysis of infected splenic M□, releasing non-viable parasites and inflating dsRed2⁺ signal in response to LdGP63_28_KO infection. Accordingly, splenic stamp smears revealed abundant lysed, non-viable amastigotes extruded from KO-infected spleens, a feature absent in parental infections, which rather showed actively proliferating amastigotes (**Fig. 3G**). Flow cytometry further confirmed the presence of these free parasites in LdGP63_28_KO infected spleen macerates, with a higher number at 12 days p.i. that dropped significantly after 24 days p.i., again hinting towards a possible parasite clearance through excess host inflammatory response (**Fig. 3I**). Thus, these results confirmed loss of LdGP63_28 yields non-persisting intracellular amastigotes due to heightened host inflammatory pyroptosis response in both *in vitro* and *in vivo L. donovani* infection.

### 5. *L. donovani* relies on GP63_31 paralogue to ensure successful host entry

After examining LdGP63_28, which revealed a limited contribution towards host attachment and initial invasion, the role of LdGP63_31 in *L. donovani* infection was further investigated. Similar to LdGP63_28_KO, knockout of *LdGP63_31* (LdGP63_31_KO) (**Fig.**□**S6A, B**) did not alter parasite morphology, growth, or motility (**Fig. S6C, D; Video S5**) and yielded metacyclic numbers comparable to the parental strain (**Fig. S6E, F**). However, two distinct parasite populations appeared within the metacyclic zone of LdGP63_31_KO (**Fig. S6E**), a feature absent in the parental and other knockout lines. Notably, attachment assay resulted in a severe defect in host attachment for LdGP63_31_KO strains (**Fig. 4A, B**). Despite this, infection assays detected intracellular LdGP63_31_KO amastigotes at□6□h□p.i., but their numbers decreased sharply by□24□h p.i. and were nearly absent by□48□h p.i. (**Fig.**□**4C**). However, in contrast to LdGP63_28_KO infections, M□ harbouring LdGP63_31_KO parasites appeared morphologically intact (**Video S6**) with pronounced dendritic extrusions during initial time of infection (**Fig. 4C**, bottom left image) without significant upregulation of apoptotic or pyroptotic markers (**Fig. 4D, E**). Time-course analyses (**Fig.**□**4F**) of intracellular infection revealed that the LdGP63_31_KO line established intracellular presence significantly earlier than the parental strain, within 0.5 h p.i., achieving a higher initial intracellular load that declined over time till 48 h p.i., producing a bell-shaped infection profile (**Fig.**□**4G**). However, despite this early establishment, LdGP63_31_KO promastigotes consistently exhibited reduced host attachment across all initial time points (**Fig.**□**4H**), consistent with previous attachment assays (**Fig.**□**4A**), indicating impaired but faster host entry resulting from defective attachment. Importantly, LdGP63_31_KO addback line complemented with a parental full-length copy of LdGP63_31 (LdGP63_31 eGFP) (**Fig.**□**S6B**) fully rescued the attachment defect observed in the knockout line (**Fig. 4I**,) while leading to a robust expression of LdGP63_31 eGFP at the host membrane (**Fig. 4J**). Furthermore, intracellular amastigote persistence was completely restored in the addback line, with substantial numbers of intracellular amastigotes detected even at 48 h p.i. (**Fig. 6K**). Interestingly, LmGP63 has previously been reported to possess a GPI anchor [29]. Consistent with this, in silico analysis predicted a higher probability of a putative GPI-anchor signal in LdGP63_31 than in LmGP63, whereas the likelihood was considerably lower for LdGP63_28 (**Fig. 4L**), supporting a role for LdGP63_31 in host membrane association.

**Figure 4:**
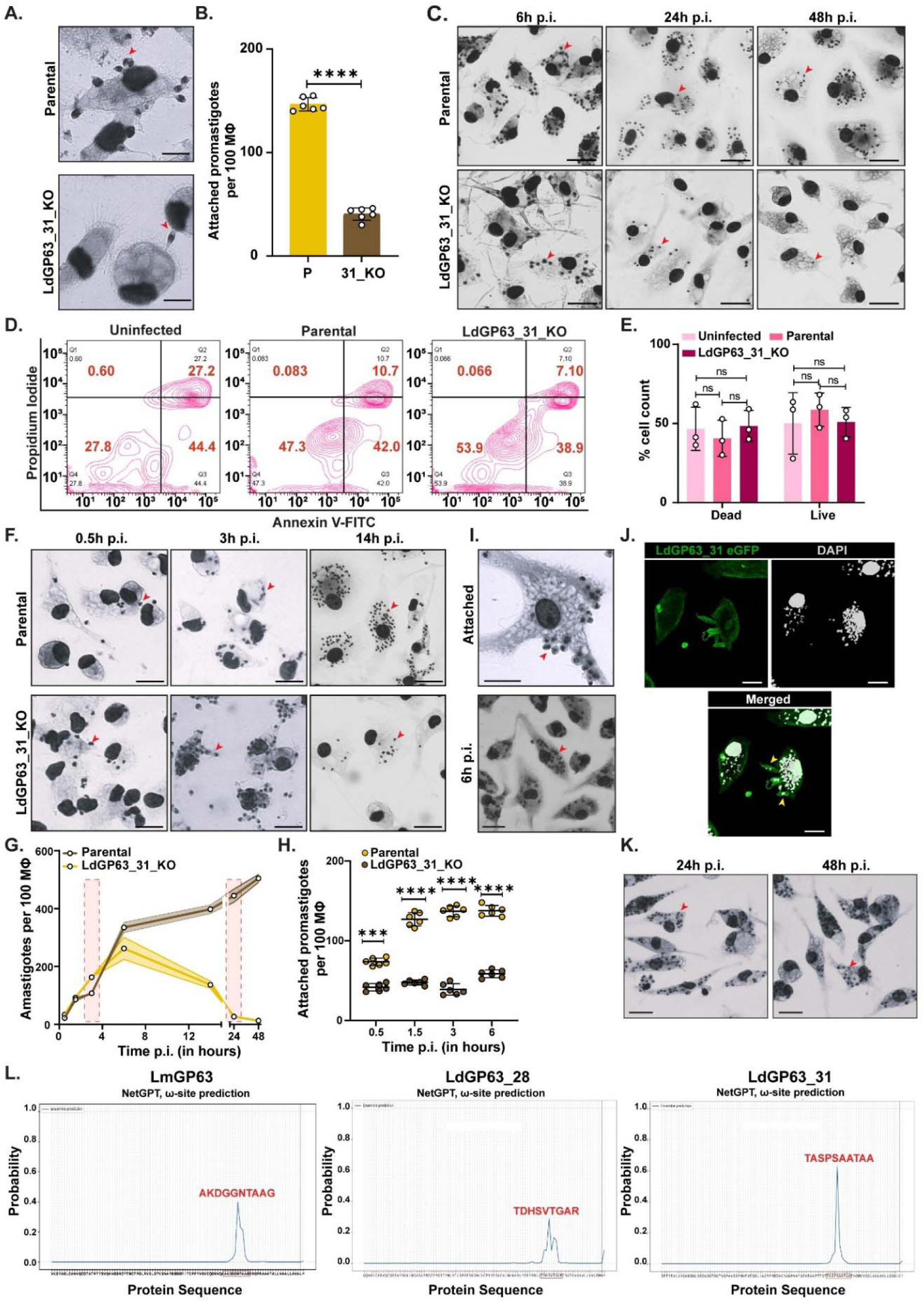
Loss of LdGP63_31 impairs the host attachment of *L. donovani* promastigotes, leading to defective host entry. A. Giemsa-stained images comparing initial attachment at 30 min p.i. Attached promastigotes are pointed by red arrows. Scale bar: 5 µm. B. Graph representing number of attached parental and LdGP63_31_KO promastigotes in 100 M□ (n = 6). C. Giemsa-stained images of murine M□ infected with parental and LdGP63_31_KO lines at 6 h, 24 h, and 48h p.i. Intracellular amastigotes are pointed by red arrows. Scale bar: 10 µm. D. Contour plot showing Propidium Iodide (PI) vs Annexin V-FITC stained cells comparing morphology of uninfected, parental, and LdGP63_31_KO infected M□ at 48 h p.i., E. Graph showing the percentage count of dead or alive M□ uninfected or infected by parental or LdGP63_31_KO parasites at 24 h p.i. (n = 3). F. Giemsa-stained images of parental and LdGP63_31_KO infected M□ at 0.5 h, 3 h, and 14 h p.i. Intracellular amastigotes are pointed by red arrows. Scale bar: 10 µm. G. Intracellular amastigote survival graph comparing the number of amastigotes from 0.5 h up to 48 h p.i. in parental and LdGP63_31_KO infected M□ (n = 3). 3 h and 24 h time points have been enclosed in transparent red boxes to show the difference between the amastigote counts. H. Scatter plot showing number of attached parental and LdGP63_31_KO parasites in 100 M□ (n = 6). I. Giemsa-stained images of M□ infected by LdGP63_31 addback lines showing attachment (top) and initial infection at 6 h p.i. (bottom). Attached parasites are marked with red arrows. Scale bar: 5 µm (top) and 10 µm (bottom). J. Representative confocal micrograph showing expression of LdGP63-31 eGFP addback promastigotes attached (yellow arrows) and infected at 6 h p.i. to murine M□. Scale bar: 5 µm. K. Giemsa-stained images of M□ infected by LdGP63_31 addback lines showing intracellular amastigotes at 24 h and 48 h p.i. Attached parasites are marked with red arrows. Scale bar: 10 µm. L. NetGPI plots of predicted GPI-sites in LmGP63, LdGP63_28, and LdGP63_31. The predicted amino acid residues of the GPI-anchor are written in red font above the predicted peak. Data are presented as mean ± standard error, with each individual point shown. Statistical analysis was performed using paired two-tailed Student’s *t*-test (B), one-way ANOVA followed by Tukey’s MCT (E), and two-way ANOVA followed by Sidak’s MCT (H): ns, non-significant (P > 0.05); *, P ≤ 0.05; **, P ≤ 0.01; ***, P ≤ 0.001; ****, P ≤ 0.0001.

### 6. Defective entry results in clearance of intracellular LdGP63_31_KO

To determine the effect of the defective entry and what resulted in the clearance of LdGP63_31 KO amastigotes without death of infected host M□, the process of internalization and morphology of internalized LdGP63_31_KO amastigotes were further investigated. SEM of M□ with attached promastigotes revealed that unlike the tight adhesive interactions of parental or LdGP63_28_KO parasites, LdGP63_31_KO promastigotes engaged in a lateral attachment and were surrounded by host M□ dendritic projections (**Fig. 5A, B**), similar to the ones observed in infection assay (**Fig. 4C**, lower left image), suggesting that impaired attachment favoured an enhanced faster phagocytic uptake rather than invasion causing premature yet defective internalization. This defective internalization of LdGP63_31_KO line was further supported by the fact that unlike parental amastigotes, LdGP63_31_KO amastigotes retained a spindle-shaped morphology with visible cytosol and flagella, indicative of defective amastigote formation (amastigogenesis) (**Fig.**□**5C**) even at 14 h p.i. TEM of M□ harbouring LdGP63_31_KO parasites (**Fig. 5D, E**) further proved this with prominent developmental defects witnessed for internalized amastigotes retaining promastigote-like structure, appearing significantly larger with apoptotic blebs and possible autophagosomes, or, in some cases, having a smaller/reduced/undeveloped dead phenotype with severely shrunken or no nuclei as compared to internalized parental amastigotes. The PV membrane surrounding internalized LdGP63_31_KO parasites, unlike parental amastigotes, also appeared to be inconsistent and ruptured in some instances. Collectively these data identify LdGP63_31 as a key mediator of host membrane attachment, required for coordinated parasite entry, essential for maintaining PV integrity, and successful amastigogenesis, thereby sustaining intracellular infection.

**Figure 5:**
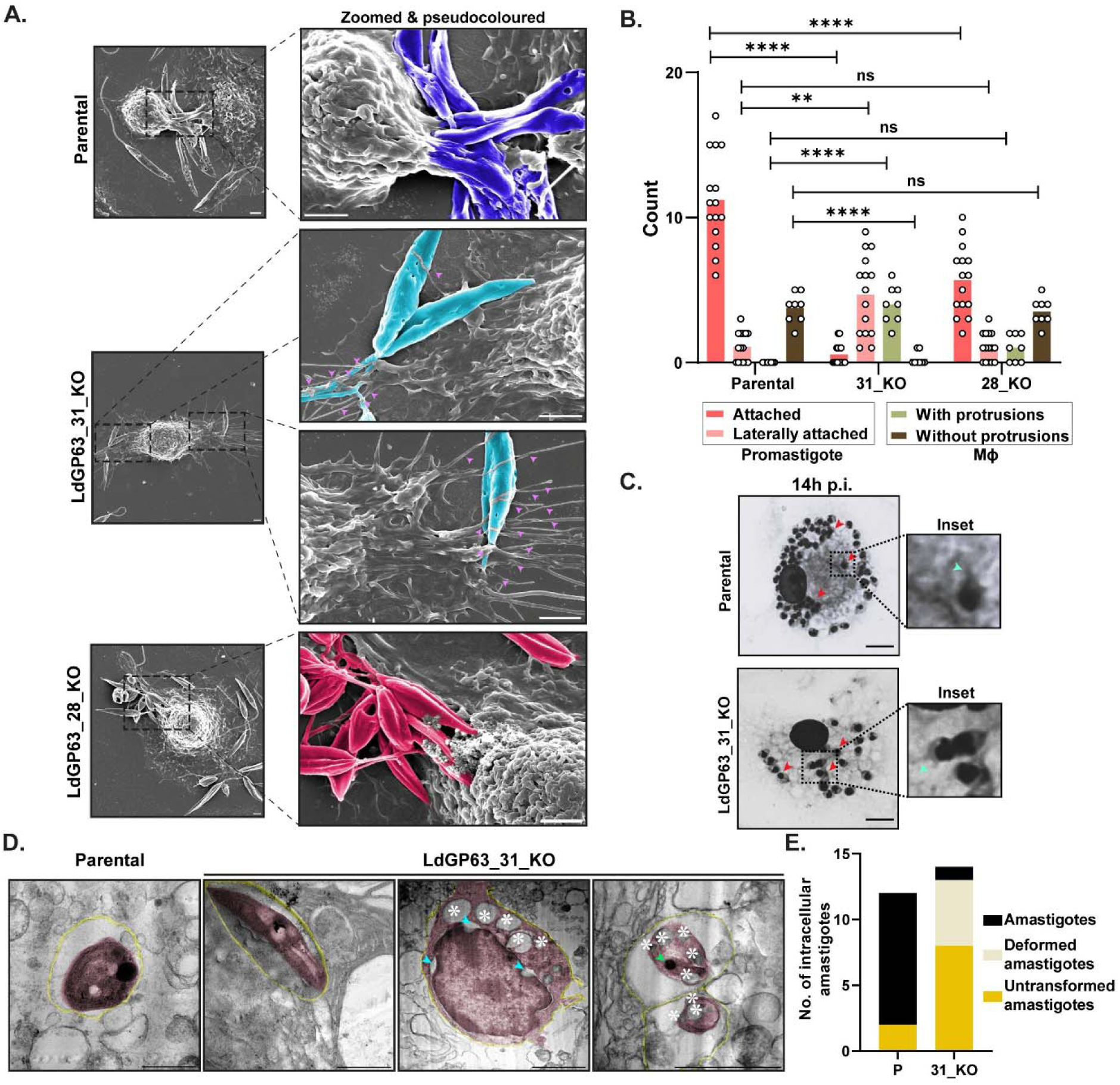
LdGP63_31_KO disrupts amastigogenesis with lateral attachment phenotype in promastigotes. A. Representative SEM images depicting the attachment of parental (blue), LdGP63_31_KO (cyan), and LdGP63_28_KO (red) promastigotes to M□ at 30 min p.i. *via* attachment assay. Dendritic projections are indicated by pink arrows. Scale bar: 5 µm. B. Bar graph representing the number of actively attached and laterally attached parental, LdGP63_31_KO, and LdGP63_28_KO promastigotes (n = 15), and M□ count with and without protrusions (n = 8), counted across different SEM images. C. Magnified (100×) image of infected M□ of parental and LdGP63_31_KO lines at 14 h p.i., with intracellular amastigotes pointed by red arrows and parasite tail by cyan arrows. Scale bar: 10 µm. D. Representative TEM images showing parental and LdGP63_31_KO intracellular amastigotes (brown) enclosed within the PV (yellow). Apoptotic blebs, shrunk nuclear gaps, and shrunk nucleus are marked by white asterisks, cyan and green arrows, respectively. Scale bar: 2 µm. E. Bar diagram showing number of amastigotes, deformed amastigotes, and untransformed amastigotes observed in TEM images of M□ infected with intracellular amastigotes (D) of parental and LdGP63_31_KO lines. Data are presented as mean ± standard error, with each individual point shown. Statistical analysis was performed using two-way ANOVA followed by Tukey’s MCT (B): ns, non-significant (P > 0.05); *, P ≤ 0.05; **, P ≤ 0.01; ***, P ≤ 0.001; ****, P ≤ 0.0001.

**Figure 6:**
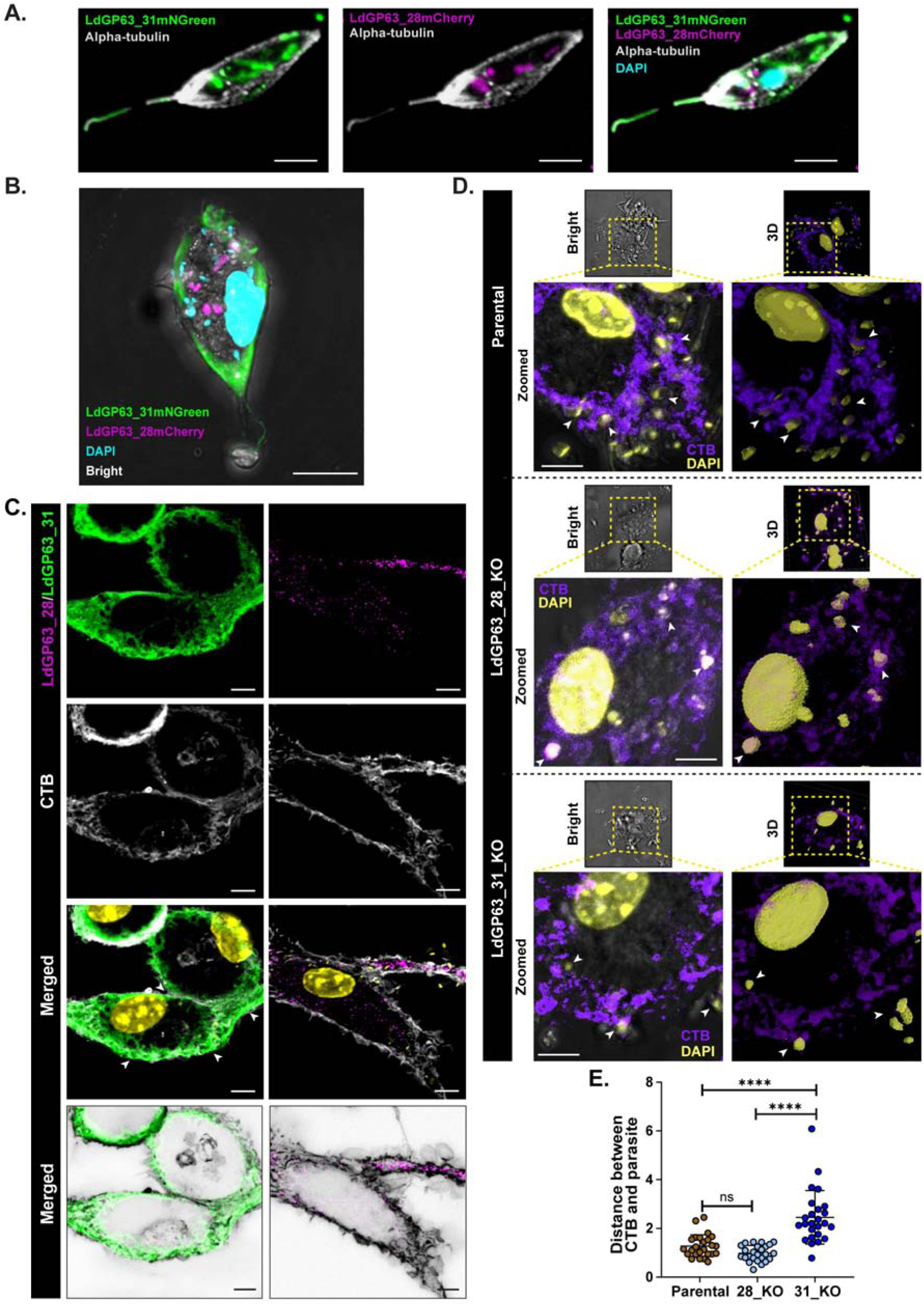
LdGP63_31 localizes in host membrane during *L. donovani* entry. A. Ultrastructure expansion of promastigotes coupled with confocal microscopy showing LdGP63_31mNeonGreen (LdGP63_31mNGreen) (green) and LdGP63_28mCherry (magenta) expression with respect to alpha-tubulin. Scale bar: 5 µm. B. Confocal micrograph showing the localization of LdGP63_31mNGreen (green) and LdGP63_28mCherry (magenta) in infected M□ 6 h p.i. Scale bar: 10 µm. C. Confocal micrographs comparing the localization of LdGP63_31mNGreen (green; LdGP63_31) and LdGP63_28mCherry (magenta; LdGP63_28) with lipid raft marker CTB (grey). Co-localized regions between LdGP63_31mNGreen and CTB is indicated by white arrow. Extreme right panel shows merged image of inverted CTB with LdGP63_28 (top) and LdGP63_31 (bottom). Scale bar: 5 µm. D. Brightfield images (top-left panels) showing attached parental, LdGP63_28_KO, and LdGP63_31_KO parasites to M□, followed by zoomed (regions marked in yellow boxes) confocal micrographs highlighting the interaction of attached parasites with CTB (violet; bottom-left panels). 3D views (generated in LAS X software) of the wide (top-right panels) and zoomed regions (bottom-right panels) are shown. White arrows indicate the parasite DAPI in close vicinity to the host lipid raft (labelled by CTB). Scale bar: 5 µm. E. Scatter plot of distance calculated between each parasite (n = 30) and its nearest M□ CTB-enriched region.

### 7. LdGP63_28 and LdGP63_31 have differential localizations

To delineate the distinct contributions of LdGP63_28 and LdGP63_31 in host entry and amastigote persistence, their subcellular localization and host interaction profiles were examined. LdGP63_31 was endogenously tagged with mNeonGreen (mNGreen) in the LdGP63_28mCherry background (**Fig. S6G, H**) and proceeded to further infect murine M□. Ultrastructure-expansion of double-tagged promastigotes (LdGP63_31mNGreen/ LdGP63_28mCherry) showed punctuated cytosolic localization of LdGP63_28 (**Fig. 6A**), consistent with previous observations (**Fig. 2D**), whereas LdGP63_31 was found to be distributed throughout the cytosol, flagellum, and cytoskeleton, partially co-localizing with alpha-tubulin (**Fig. 6A**). Following M□ infection, LdGP63_28mCherry predominantly localized within the host cytosol, while LdGP63_31mNGreen was enriched in the M□ membrane (**Fig. 6B**). Importantly, localization of both endogenously tagged LdGP63_31mNGreen and LdGP63_28mCherry in promastigotes and infected M□ closely mirrored LdGP63_31 and LdGP63_28 localization observed upon second-copy eGFP expression in the LdGP63_31 and LdGP63_28 addback line, further validating the previously observed distribution (**Fig. 4J)**. Co-localization with Cholera toxin B (CTB) further demonstrated that LdGP63_31 was significantly more associated with lipid raft microdomains on host membrane than LdGP63_28 (**Fig. 6C**). Consistently, enrichment of host lipid rafts encircling the parasites during their entry was significantly higher in case of parental, while they appeared to be perfectly co-localizing with LdGP63_28_KO parasites, but was selectively diminished during interactions with LdGP63_31_KO parasites. (**Fig. 6D, Video S7**). Additionally, LdGP63_31_KO parasites remained mostly farther away from the CTB-enriched regions (**Fig. 6F**), confirming a specific and essential role for LdGP63_31 in host membrane engagement during promastigote entry.

### 8. Localization specificity of LdGP63_28 and LdGP63_31 is complemented by their differential substrate specificities

Next, we investigated whether the distinct localization of LdGP63_28 and LdGP63_31 in infected host can be explained by their differential substrate preferences during *L. donovani* infection. Their interactions with five reported LmGP63 substrates (mTOR, C3b, c-Jun, MARCKS, and SHP-1) [15, 31, 33–37] of defined host localization were examined (**Table 1**). However, the docking of LmGP63, LdGP63_28, and LdGP63_31 with substrates yielded comparable values, making it difficult to infer distinct substrate specificities of the two proteases (**Fig. 7A-F, Fig. S7A-D).** Furthermore, 100 ns molecular dynamics simulations of both proteases in complex with mTOR, C3b, and c-Jun, selected on the basis of their distinct and well-defined subcellular localizations in the *L. donovani* infected host, also showed stable interactions, as indicated by root mean square deviation (RMSD) (**Fig. S7E**), root mean square fluctuation (RMSF) (**Fig. S7F**), radius of gyration (Rg) (**Fig. S7G**), and hydrogen bonding (**Fig. S7H**) values (**Table S3**).

**Figure 7:**
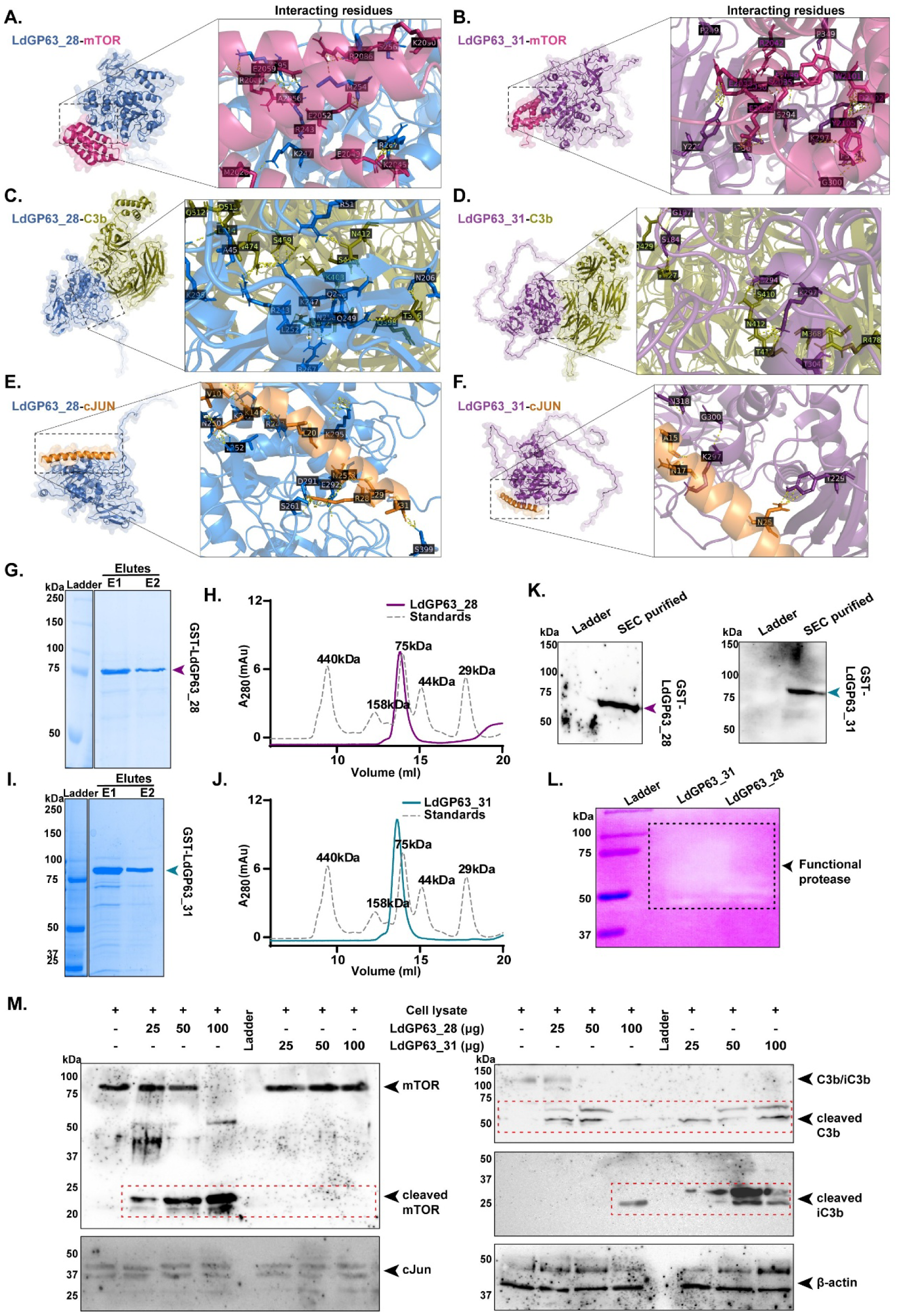
LdGP63_28 and LdGP63_31 have preferential substrate specificities. A-F. Interactive enzyme-substrate diagrams of LdGP63_28 and LdGP63_31 with mTOR (A, B), C3b (C, D) and c-Jun (E, F) highlighting the different interacting amino acid residues with their respective positions (right-side panel). G-J. Coomassie blue stained SDS-PAGE gel showing glutathione affinity purified fractions of N-terminal GST-tagged LdGP63_28 (G) and LdGP63_31 (I), followed by their size exclusion chromatrogram (SEC) peaks (H and J for LdGP63_28 and LdGP63_31, respectively). The dotted peaks in H and I correspond to the standard protein peaks. K. Western blot of the SEC purified GST-LdGP63_28 (left) and GST-LdGP63_31 (right) fractions. L. Zymogram of the SEC purified proteases visualized on non-denaturing 1.2 mg/ml gelatin-containing PAGE gel. The white clear band enclosed in black dotted box indicates protease activity. M. Western blots analysis of mTOR, C3b, and c-Jun of M□ lysate in the absence and presence of recombinant LdGP63_28 and LdGP63_31 at varying concentrations of 25 µg, 50 µg, and 100 µg. Lanes pertaining to cleaved bands are enclosed in red-dotted rectangles. β-actin was used as the loading control.

**Table 1:**
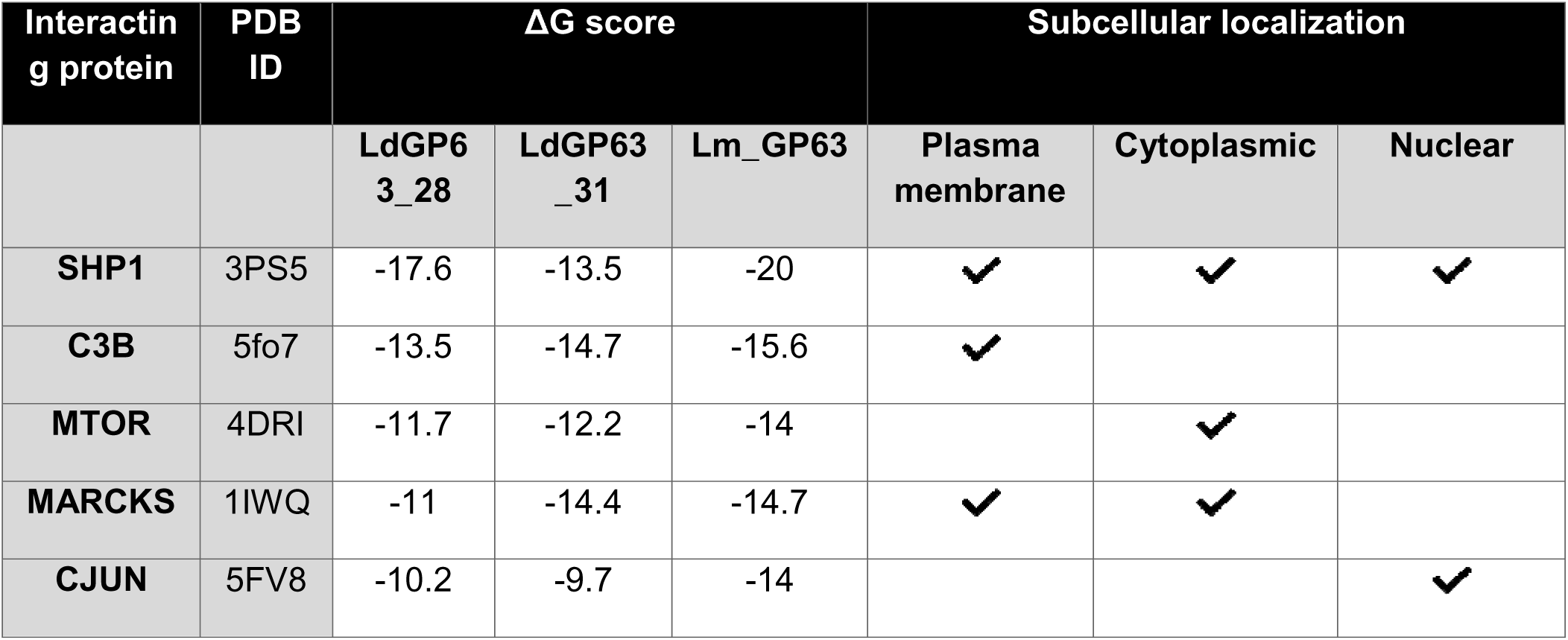
Docking scores of some known Lm_GP63 substrates with LdGP63_28 and LdGP63_31, indicating their subcellular localizations.

| Interactin<br>g protein | PDB<br>ID | $\Delta G$ score | | | Subcellular localization | | |
| --- | --- | --- | --- | --- | --- | --- | --- |
|  |  | LdGP6<br>3_28 | LdGP63<br>_31 | Lm_GP63 | Plasma<br>membrane | Cytoplasmic | Nuclear |
| **SHP1** | 3PS5 | -17.6 | -13.5 | -20 | ✓ | ✓ | ✓ |
| **C3B** | 5fo7 | -13.5 | -14.7 | -15.6 | ✓ |  |  |
| **MTOR** | 4DRI | -11.7 | -12.2 | -14 |  | ✓ |  |
| **MARCKS** | 1IWQ | -11 | -14.4 | -14.7 | ✓ | ✓ |  |
| **CJUN** | 5FV8 | -10.2 | -9.7 | -14 |  |  | ✓ |

Since computational analyses to identify preferable substrates of LdGP63_28 and LdGP63_31 in *L. donovani* infected M□ remained inconclusive, enzymatic (proteolytic) assays were performed. Purified recombinant LdGP63_28 (**Fig. 7G, H**) and LdGP63_31 (**Fig. 7I, J**) were validated by immunoblotting (**Fig. 7K**). Zymograms demonstrated that both enzymes were catalytically active (**Fig. 7L**). *In vitro* cleavage assays with increasing concentration of purified proteases confirmed functional divergence, although with different outcomes from the docking scores, where mTOR showed selective cleavage by LdGP63_28, while LdGP63_31 specifically processed the membrane secreted C3b, generating both cleaved C3b and iC3b fragments (**Fig. 7M**). Although LdGP63_28 also cleaved C3b, this reaction mostly yielded C3b and iC3b only at higher concentration (100 µg), consistent with its limited role in host entry and a primary function in intracellular amastigote maintenance. Neither protease cleaved c-Jun or β-actin. Together, these findings established that LdGP63_31 primarily mediates host attachment through lipid raft-associated substrates, whereas LdGP63_28 predominantly supports intracellular survival by targeting cytosolic host factors.

### 9. LdGP63_28 plays a more prominent role in egress and re-invasion of intracellular *L. donovani* amastigotes

To determine the relative contributions of GP63 paralogues during amastigote infection, axenic and intracellular amastigotes from the double-tagged LdGP63_28mCherry–LdGP63_31mNGreen line were purified and analysed by confocal microscopy. Both proteases were expressed in axenic as well as intracellular amastigotes (**Fig. 8A**). Infection assays using these purified amastigotes revealed localization patterns analogous to those observed during promastigote infection with LdGP63_28 retaining a punctate cytosolic distribution within host, whereas LdGP63_31 was predominantly enriched at the host plasma membrane (**Fig. 8B**), suggesting that both proteases may participate during amastigote re-infection. Next, axenic amastigotes generated from parental, LdGP63_28_KO, and LdGP63_31_KO lines were used to infect M□ (**Fig. 8C**). Notably, LdGP63_31_KO parasites exhibited reduced amastigogenesis with structural defects (**Video**□**S8**), as we have previously observed for intracellular amastigotes with respect to LdGP63_31_KO infection (**Fig. 5E**). SEM imaging confirmed these morphological differences with LdGP63_31_KO axenic amastigotes appearing thicker, longer, and often flagellated, while LdGP63_28_KO forms were mildly elongated but largely flagella-deficient (**Fig. 8D, E**). However, despite the compromised amastigogenesis, LdGP63_31_KO axenic amastigotes still retained significant infection capacity, whereas LdGP63_28_KO axenic amastigotes completely failed to establish infection at 24□h□p.i., supporting a stronger role for LdGP63_28 in amastigote invasion (**Fig.**□**8F, G**). Re-infection assays with syringe-lysed intracellular amastigotes (**Fig. 8H**) from parental, LdGP63_28_KO, and LdGP63_31_KO lines into fresh M□, revealed, in contrast to axenic infections, marked loss of re-infection ability in both KO forms (**Fig.**□**8I, J**). Therefore, while both proteases may contribute, LdGP63_28 plays a more prominent role in amastigote re-infection. This difference in infectivity between axenic and intracellular amastigotes of LdGP63_31_KO could also result from a loss of viability of isolated intracellular amastigotes as LdGP63_31_KO amastigotes fail to sustain within host M□, and get cleared within a very short period of time as compared to intracellular LdGP63_28_KO amastigotes, compromising their fitness to re-infect.

**Figure 8:**
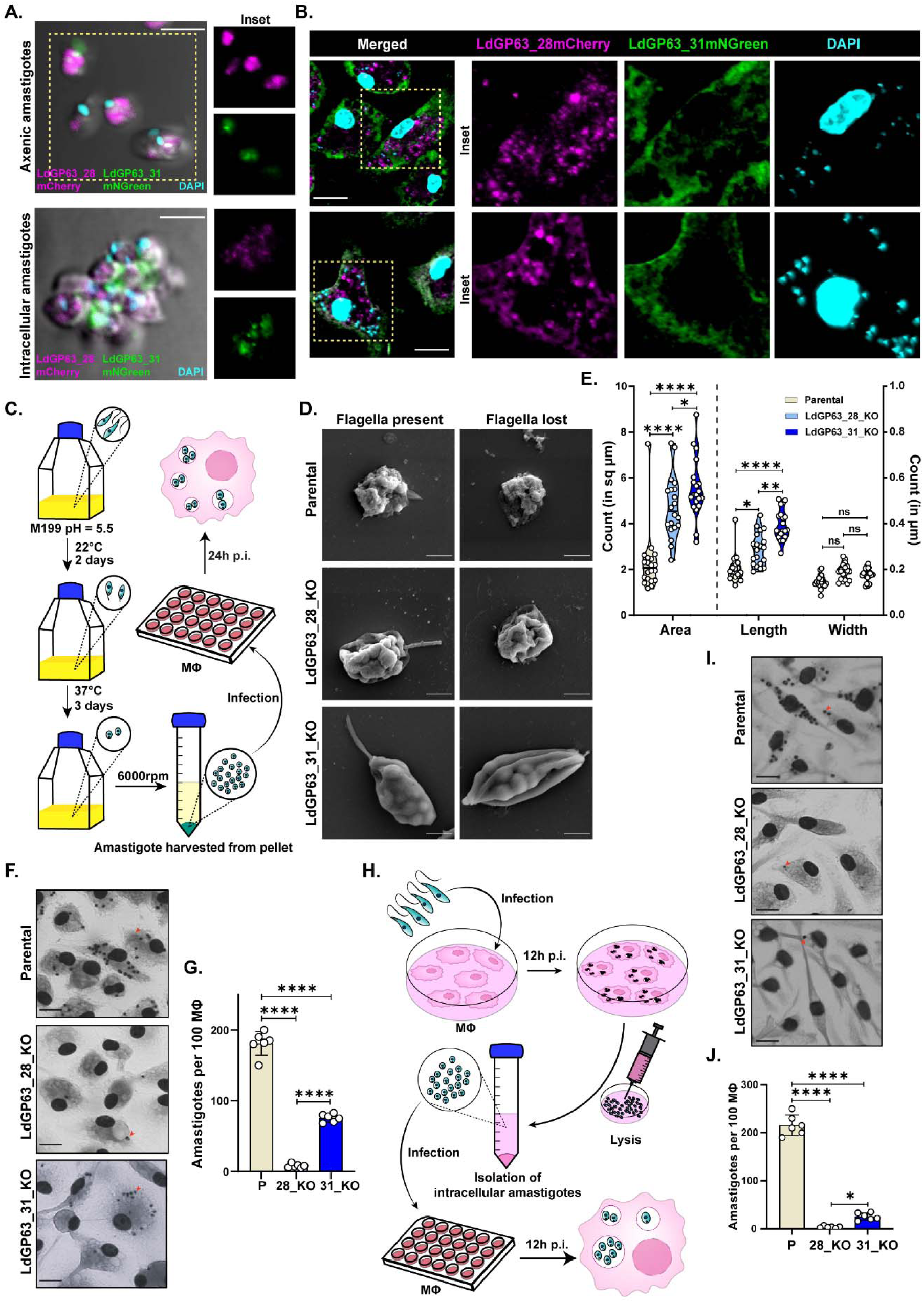
LdGP63_28 plays a critical role in amastigote re-infection. A. Axenic and intracellular amastigotes of LdGP63_28mCherry-LdGP63_31mNGreen showing expression of LdGP63_28 (magenta) and LdGP63_31 (green). Scale bar: 2 µm. B. Localization of LdGP63_28 (magenta) and LdGP63_31 (green) in axenic (top panel) and intracellular (bottom panel) amastigotes infected M□. Scale bar: 2 µm. C. Schematic methodology for generation of axenic amastigotes for further M□ infection assay. D. SEM micrographs comparing axenic amastigotes of parental, LdGP63_28_KO, and LdGP63_31_KO lines with stumped flagella (left panel) and no flagella (right panel). Scale bar: 1 µm. E. Violin plot representing the area (sq µm), length (µm), and width (µm) of axenic amastigotes. Each dot represents data from one axenic amastigote (n = 20). F. Giemsa-stained images of parental, LdGP63_28_KO, and LdGP63_31_KO axenic amastigote infected M□ at 24 h p.i. Intracellular amastigotes are pointed by red arrows. Scale bar: 10 µm. G. Representative bar graph showing infectivity of parental, LdGP63_28_KO, and LdGP63_31_KO axenic amastigotes (n = 6). H. Schematic methodology for isolating intracellular amastigotes from infected M□ and reinfection of fresh M□. I. Giemsa-stained images of M□ infected with intracellular amastigotes (red arrows). Scale bar: 10 µm. J. Representative bar graph showing infectivity of parental, LdGP63_28_KO, and LdGP63_31_KO intracellular amastigotes (n = 6). Data are presented as mean ± standard error, with each individual point shown. Statistical analysis was performed using two-way ANOVA followed by Tukey’s MCT (E) and one-way ANOVA followed by Tukey’s MCT (G, J): ns, non-significant (P > 0.05); *, P ≤ 0.05; **, P ≤ 0.01; ***, P ≤ 0.001; ****, P ≤ 0.0001.

## DISCUSSION

*Leishmania* spp. cause either localized dermal disease or systemic visceral infection, reflecting fundamentally different host microenvironments and immune pressure challenges [3]. This adaptability is driven by an unusually plastic genome characterized by mosaic aneuploidy, copy-number variation, extrachromosomal amplification, and recombination, generating rapid phenotypic diversity that supports survival across disparate host niches [38, 39]. Additionally, with hybridization and parasexual processes [40–42], this plasticity is thought to underpin specific tissue adaptation, exemplified by visceral dissemination of *L. donovani* versus dermal restriction of *L. major* [43, 44]. However, the molecular determinants translating genomic flexibility into stage and tissue-specific manifestation have remained poorly defined.

While GP63, a zinc metalloprotease, is a canonical virulence factor in cutaneous leishmaniasis [14], its role in its visceral disease has remained ambiguous. The *gp63* locus is among the most polymorphic regions of the *Leishmania* genome, with marked differences in paralogue number, chromosomal distribution, and ploidy between dermal and visceral species [16, 19, 20]. This study highlights that *L. donovani gp63* paralogues have not simply been retained or expanded, but have evolved independently into functionally specialized modules that collectively enable visceral colonization. This specialization contrasts with the architecture of dermatropic species and provides a mechanistic link between genome plasticity and organ-specific pathogenesis.

The initial phylogenetic analysis resolved *L. donovani* GP63 into two major clades encoded on chr-31 (*LdGP63_31*) and chr-28 (*LdGP63_28*) (**Fig. 1B**), consistent with their functional divergence. In contrast to *L. major*, where the chr-10 harbors four copies of GP63 and act as the dominant virulence determinant [45], both copies of *L. donovani* chr-10 GP63 (*LdGP63_10.51* and *LdGP63_10.52*) were truncated, catalytically inactive, and dispensable for parasite growth and host infection (**Fig. S3**), highlighting lineage-specific reprogramming of the GP63 repertoire. Interestingly, our result (**Fig. 1B**) also pointed out that unlike *LdGP63_10.51* and *LdGP63_10.52, L. infantum*, which also causes visceral infection, has retained multiple copies of GP63 in chr-10 [46]. However, there are no reports characterising their functional activity and impact in visceral infection. Moreover, in contrast to earlier reports which highlighted deletion of *gp63* impairs host infection in *L. major* [25] , a recent report has highlighted deletion of all four copies of chr-10 GP63 in *L. major* affects intracellular growth [29], indicating that unresolved complexities in GP63-mediated dermal adaptation may still exist. Deeper evaluation into amino acid residues within the conserved motifs of LmGP63, LdGP63_28 and LdGP63_31 uncovered amino acid substitutions among all three proteases (**Fig. 1C**), which could probably have led to a functional diversification and eventual disease outcome between *L. major* and *L. donovani* in due course of evolution.

Unlike chr-10 GP63s, LdGP63_28 retained an intact active site and localized to cytosolic vesicles in promastigotes and infected macrophages (**Fig. 2**). Although loss of LdGP63_28 had only minor effects on parasite attachment and host entry, LdGP63_28 proved essential for intracellular amastigote persistence. Loss of LdGP63_28 resulted in failure to suppress host inflammatory responses and progressive parasite clearance (**Fig. 2**). Mechanistically, LdGP63_28-deficiency biased infected M□ towards inflammasome-driven pyroptosis, marked by Gasdermin-E cleavage and Caspase-3 activation (**Fig. 3, Fig. S5**). While suppression of pyroptosis has also been implicated in host survival of dermal *Leishmania* spp. [47–49], our data position LdGP63_28 as a critical regulator of inflammatory host cell death that stabilizes the intracellular niche required for visceral *L. donovani* amastigote replication, a function that may extend to dermal amastigotes.

In contrast, LdGP63_31 functioned at the initial stages of infection. LdGP63_31_KO promastigotes exhibited profound defects in M□ attachment and coordinated entry (**Fig. 4**, **Fig. 5**), undergoing aberrant faster uptake *via* lateral dendritic interactions that failed to support productive biogenesis of parasitophorous vacuole (PV). Internalized LdGP63_31_KO parasites were unable to complete amastigogenesis retaining promastigote-like morphology, or displayed degenerative features, culminating in rapid clearance without death of infected M□ (**Fig. 5D, E**). Consistent with this role, GPI-anchored LdGP63_31 was selectively enriched in host lipid rafts during initial parasite-M□ contact (**Fig. 6C**). Loss of LdGP63_31 hampered the interaction between the parasites with the host lipid raft, implicating LdGP63_31 in membrane remodeling required for productive entry and PV formation (**Fig. 6D**). Interestingly, LdGP63_28_KO resulted in enhanced superimposition of lipid rafts on the attached parasites, contrary to parental infection where the lipid rafts oriented themselves encircling the parasites, highlighting that the loss of LdGP63_28 must have led to an altered host lipid raft related signalling pathways (**Fig. 6D**) thereby differentially regulating Ld persistence.

Substrate profiling further reinforced this functional partitioning. LdGP63_31 preferentially processed host membrane-associated complement components such as C3b to generate iC3b, facilitating immune evasion and entry, whereas LdGP63_28 targeted cytosolic regulators of inflammatory signalling (**Fig. 7**, **Fig. S7**, **Table 1**).

Stage-specific infection assays further reinforced these distinct roles, while LdGP63_28_KO promastigotes differentiated efficiently into axenic amastigotes, their infectivity was markedly reduced, underscoring the essential role of LdGP63_28 during amastigote infection. In contrast, LdGP63_31_KO parasites showed impaired amastigogenesis, yet retained partial infectivity under axenic conditions (**Fig. 8**). Emergence of two distinct metacyclic populations in LdGP63_31-deficient promastigotes (**Fig. S6E**) and its impaired axenic amastigogenesis (**Fig. 8D**) further support its role as an active protease involved in membrane and cytoskeletal remodelling, reinforcing division of labour between these two paralogues through independent evolution to support visceral infection through stage- and compartment-specific specialization.

## MATERIALS AND METHODS

### Homology searches and phylogenetic analysis

The sequences of identified *L. donovani* GP63 proteins (LdBPK_100510.1, LdBPK_100520.1, LdBPK_280600.1, and LdBPK_312040.1) were retrieved from TriTrypDB release 68 [28] . Further, GP63 in *L. donovani* and other species (**Table S1**) were identified in TriTrypDB or by blastp and tblastn searches v2.12.0+ [50] against genome-derived proteomes and genomes, respectively, from TriTrypDB or a previous publication [51]. The identified sequences were aligned by MAFFT 7.458 [52] using the L-INS-i algorithm. The ambiguously aligned positions were removed by BMGE v1.12 [53]. Maximum-likelihood phylogenetic analysis was performed in IQ-TREE v2.3.5 [54] using the PMSF method [55] with the LG+C60+G4 model, the guide tree inferred with the LG+G model, 1,000 replicates for ultrafast bootstraps [56] and Shimodaira-Hasegawa approximate likelihood ratio test (SH-aLRT) [57], and a maximum of 5,000 iterations.

### GP63 alignment, sequence annotation and predictions

The LdGP63 proteins were aligned with the already available *L. major* GP63 (LmjF.10.0460, LmGP63) sequence using the MUSCLE tool [58] at EMBL-EBI [59]. The alignment file was annotated in ESPript v3.0 beginner mode [60]. LmGP63 available from Protein Data Bank (PDB; 1LML) was uploaded as the secondary structure, and labels were set to α1, β1, α2, β2. Considering the crystal structure of LmGP63 as the reference [29], the N-terminal, central, and C-terminal domains were labelled highlighting the conserved regions (PAVGVINIPA, KAREQYGC and QDELMAP) and HEXXH active site. Schematic diagrams highlighting the domains and sites was generated using DOG v2.0 - Protein Domain Structure Visualization [61]. Signal peptide regions were predicted using DTU/SignalP-6 [62]. NetGPI v1.1 [63] tool was used to predict the presence GPI-anchors in the proteases.

### Homology modelling and visualization

The crystal structure of LmGP63 was retrieved from PDB and 3D structures of GP63_28 and LdGP63_31 were modelled using AlphaFold v2 [64] and visualized with PyMOL v2.5 (The PyMOL Molecular Graphics System, Version 2.5 Schrödinger, LLC.). TM-scores and RMSD scores were calculated by TM-Align tool [65] and in PyMOL using the “Align” plugin in “Action” menu, respectively. The stereochemical qualities of LdGP63_28 and LdGP63_31 models were validated by analysing Ramachandran plots in PROCHECK [66] *via* the SAVES v6.1 (Structure Analysis and Validation Server, UCLA-DOE Institute). Validating the positions of the amino acid residues, labelling them, and assigning colours to the structures were all done in PyMOL.

### Protein-protein docking and ΔG calculation

Docking was performed in HADDOCK v2.4 web server [67] under default parameters . Each run involved one protease (LmGP63, LdGP63_28, or LdGP63_31) with a reported substrate of LmGP63: complement C3b, myristoylated alanine-rich protein kinase C substrate (MARCKS), mechanistic target of rapamycin (mTOR), Src homology region 2 domain-containing phosphatase-1 (SHP-1), and transcription factor jun (c-Jun) [14, 15, 25, 31, 34, 37], PDB IDs of which are listed in **Table 1**. The resulting docked complexes (four for each interaction) were obtained and their respective binding affinities (ΔG) were calculated using the PRODIGY web server [68]. Lowest ΔG values for each interaction were interpreted as stronger protein-protein interactions than the others and were used for further validation. For structure visualization and interpretation, PyMOL was used where each protein chain was coloured individually to distinguish binding partners in the docked complexes. LIGPLOT+ software v2.3 [69] was utilized to identify the interacting residues between the proteases and the substrates, which were then annotated in PyMOL for representation.

### Molecular dynamics (MD) simulation

MD simulations of the protein–protein (PPI) complex were performed using GROMACS v2020.4 with the AMBER99SB force field [70] and an explicit TIP3P water model. The docked PPI complex was placed in a cubic simulation box (8.85499 × 8.85499 × 8.85499 nm³), maintaining minimum 10 Å buffer between the solute and box edges. After solvation, the system was neutralised, and additional ions were added to reach a physiological salt concentration (0.15 M NaCl) using GROMACS gmx genion (-neutral -conc 0.15), replacing water molecules with Na⁺and Cl⁻ ions. Energy minimisation was performed using the steepest-descent algorithm for 50,000 steps until the maximum force converged to below 500 kJ/mol/nm.

After minimization, position restraints were applied to the protein complex, and the system was equilibrated under an NVT ensemble for 100 ps at 300 K using the V-rescale thermostat with a 2 fs time step. This was followed by NPT equilibration for 100 ps using the Parrinello–Rahman barostat with isotropic pressure coupling, maintaining a temperature of 300 K and a pressure of 1 bar. Subsequently, production MD was performed for 100 ns using a leapfrog integrator with a 2 fs time step, and the coordinates were saved every 10 ps. The structural stability and compactness of the complex were evaluated in terms of the root mean square deviation (RMSD), root mean square fluctuation (RMSF), hydrogen bonds, and radius of gyration (Rg) using GROMACS tools (gmx rms, gmx rmsf, gmx hbond, and gmx gyrate).

### Maintenance of parasite strains

Promastigotes of *L. donovani* strain MHOM/IN/09/BHU575/0 and all other mutant strains were cultured at 22°C in M199 medium (Sigma-Aldrich) supplemented with 5% NuSera (Himedia) [71]. The appropriate selection drugs (SRL) were added to the medium at the following concentrations: 200 µg/ml hygromycin B, 200 µg/ml blasticidin dihydrochloride, 200 µg/ml G418, and 200 µg/ml puromycin dihydrochloride.

### qRT-PCR analysis

For gene expression analysis, 2 × 10^8^ log-phase promastigotes, or murine M□ infected with *L. donovani* amastigotes were harvested and RNA was extracted according to the RNeasy Mini kit protocol (Qiagen). To synthesize complementary DNA (cDNA), 2 μg of RNA was reverse-transcribed using the ProtoScript^®^ II First Strand cDNA Synthesis Kit (NEB). Quantitative RT-PCR was then performed with 200 ng of cDNA per sample using SYBR Green Master Mix (Applied Biosystems). Data were analysed by 2^−ΔΔCT^ method through normalization with GAPDH or beta-actin. The thermal cycling conditions were 94°C for 10 min, followed by 40 cycles at 94°C for 15 sec and 60°C for 1 min. Primers used for all RT-PCR reactions are listed in **Table S4**, and the codes are as follows: LdBPK_100510.1 (P1, P2), LdBPK_100520.1 (P3, P4), LdBPK_280600.1 (P5, P6), LdBPK_312040.1 (P7, P8), gapdh (P9, P10), caspase 1 (P41, P42), caspase 3 (P43, P44), nlrp3 (P45, P46), caspase 2 (P47, P48), caspase 7 (P49, P50), nlrc4 (P51, P52), aim2 (P53, P54), nlrp1 (P55, P56), β-actin (P57, P58).

### Generation of endogenously tagged and knockout para site strains

To generate *L. donovani* cell lines that stably express Cas9 and T7 polymerase [72], pTB007 plasmid was transfected into promastigotes and Cas9 expression was confirmed by western blotting (**Fig. S2A**). This Ld-Cas9 was used as the parental strain in all experiments that followed. The primers for tagging and knockout were retrieved from the LeishGEdit website [73] and are listed in **Table S4**.

pPLOTv1 puro-mCherry-puro [72], pPLOTv1 blast-mNeonGreen-blast [72], and pPLOTv1 puro-eGFP-puro (lab cloned) were used to endogenously tag the C-terminals of LdGP63_28, LdGP63_31, and LdGP63_10.51, respectively. Briefly, the 3′ sgRNA template was amplified using the Downstream_sgRNA primer (LdGP63_28: P19, LdGP63_31: P28) and G00 primer (P29), whereas mCherry-puro, mNeonGreen-blast, and eGFP-puro cassettes were amplified from their respective plasmids using the Downstream forward primers and Downstream reverse primers (LdGP63_28: P17, P18; LdGP63_31: P26, P27, respectively). PCR reactions were performed using Q5 High-Fidelity DNA Polymerase (New England Biolabs). The thermal cycling conditions for sgRNA were 98°C for 30 sec, followed by 40 cycles at 98°C for 10 sec and 60°C for 30 sec, 72°C for 15 sec, and a final elongation at 72°C for 10 min; for the cassettes, 94°C for 5 min, followed by 40 cycles at 94°C for 30 sec and 66°C for 30 sec, 72°C for 3 min, and a final elongation at 72°C for 10 min. Amplified DNA was assessed by 2% and 1% agarose gel electrophoresis for sgRNA and for the cassettes, respectively.

To generate knockout lines, the 3′ sgRNA and 5’ sgRNA templates were amplified using the Downstream_sgRNA (LdGP63_10.51: P14, LdGP63_28: P19; LdGP63_10.52: P23, LdGP63_31: P28) and G00, and Upstream_sgRNA (LdGP63_10.51: P11; LdGP63_28: P15, LdGP63_10.52: P20, LdGP63_31: P24) and G00 primers, respectively. Puromycin and blasticidin resistance cassettes were amplified from pTpuro v1 and pTblast v1 plasmids [72], using Upstream forward primers and Downstream reverse primers (LdGP63_10.51: P12, P13, LdGP63_28: P16, P17; LdGP63_10.52: P21, 22, LdGP63_31: P26, P27, respectively), with Q5 High-Fidelity DNA Polymerase (New England Biolabs) and were together transfected into Ld-Cas9.

### Parasite transfections

For all transfections, 1 × 10^10^ parasites were mixed in a 2 mm cuvette with the relevant plasmid (20 µg) or amplified DNA (approx. 60 µl) in about 340 µl of 1× electroporation buffer (21 mM HEPES, 137 mM NaCl, 5 mM KCl, 0.7 mM NaH2PO4, 6 mM glucose), and electroporated using a BTX Electroporation system with 2 pulses at 450 V, 500 µF, and 25 Ω [74]. Electroporated cells were immediately kept on ice for 5 min and then transferred into M199 medium (substituted with 10% FCS) and left to recover overnight before adding selection drugs. Drug-resistant mutants appeared healthy about 20 days after transfection and were confirmed about 30-40 days later by PCR followed by agarose gel electrophoresis.

### DNA extraction and PCR confirmation

For PCR verification of genetically modified parasite strains and for cloning genes into plasmids, DNA was extracted using the DNAsure mini kit (Qiagen) and PCRs were performed using GoTaq DNA Polymerase (Promega) with cycling conditions as follows: 95°C for 2 min, followed by 35 cycles at 95°C for 1 min, 56°C for 1 min, 72°C for 2 min, and a final elongation at 72°C for 5 min. Amplified DNA was assessed by agarose gel electrophoresis. Primers used for the PCR are listed in **Table S4** and are as follows: for amplification of endogenous cassettes- P33, P34; for confirmation of knockouts- LdGP63_10.51 (P1, P2), LdGP63_10.52 (P3, P4), LdGP63_28 (P29, P30) and LdGP63_31 (P31, P32).

### Promastigote growth curve analysis

The promastigote growth curve was determined by following a previously published protocol [75]. In brief, approximately 1 × 10^5^ wild type and KO promastigotes were cultured in 5% NuSera (Himedia) supplemented M199 media and counted in a Neubauer chamber daily for up to 7 days. Data points were plotted on Prism v8 (GraphPad) to generate the curve.

### Scanning electron microscopy (SEM)

The promastigotes or M□ adhered in the polylysine-coated coverslips were further processed for SEM imaging [76]. Briefly, log-stage promastigotes or M□ were washed with PBS followed by incubation for 1 h in 5% OsO_4_ under a fume hood. Then, increasing ethanol gradient dehydration with 70%, 80%, 90%, and finally 100%, 10 min twice for each step, was performed. Finally, ethanol was removed, and dried and stored in a vacuum desiccator till imaging. During imaging, sections were sputtered by gold using Quorum Q150R ES and subjected to Field Emission-Merlin, Zeiss Sigma 300 VP and JEOL scanning electron microscopy units.

### Quantification of metacyclic promastigotes

Stationary phase *L. donovani* promastigotes were washed and resuspended in 1× PBS before being subjected to flow cytometry by following an already established protocol [77]. *L. donovani* populations were checked with forward scatter (FSC) versus side scatter (SSC) and dot plot was generated to represent the acquisition of 10,000 events using a FACS Aria II system. Histogram to compare between parental and knockout strains, each with three replicates, was plotted in Prism.

### Cell culture and infection

Peritoneal M□ were harvested from BALB/c mice as mentioned in an already published protocol [78], with slight modifications. After 48 h of mice intraperitoneal injection with 4% starch, M□ were isolated and plated on 35-mm petri dishes or on sterile coverslips in 24-well plates at a density of 2 × 10^6^ or 1.5 × 10^5^, respectively, and allowed to grow in RPMI 1640 medium (Gibco) supplemented with 10% NuSera (Himedia) and 1% (v/v) Penicillin-Streptomycin solution (Gibco). The cells were allowed to adhere for 48 h at 37°C in 5% CO_2_. Then, M□ were infected with metacyclic-stage sorted (Beckman Coulter Cytoflex SRT) parental or mutant strains of *L. donovani* promastigotes at 1:10 ratio (10 parasites per 1 M□), as described previously [76]. Following a 6 h incubation, the M□ were washed with 1× PBS and subsequently cultured for specific time points up to 24 h or 48 h as per experimental requirements. Coverslips were retrieved at necessary time points, rinsed with 1× PBS, fixed using ice-cold methanol or 2% paraformaldehyde, and either stained with Giemsa or proceeded for immunofluorescence staining. The prepared coverslips were then affixed onto glass slides and examined utilizing either light microscope (Olympus) or confocal microscope (see below). Parasite quantification was conducted by counting number of amastigotes per 100 M□.

### Promastigote attachment assay

The attachment of promastigotes to the murine M□ was studied using a previously reported protocol [30]. Briefly, 1.5 × 10^5^ parasites were added to M□ already seeded on a 24-well plate, by resuspending in RPMI 1640 medium (Gibco), followed centrifuging at 720 *g* at 4°C for 5 min. The plate was then incubated at 4°C for 5 min, and cells were fixed with ice-cold methanol. This was followed by Giemsa staining and the coverslips, affixed onto glass slides, were examined under Olympus light microscope. Attached parasites per 100 M□ were counted and plotted.

### Lipid raft labelling and calculations

To label the lipid rafts, stationary-phase parental, LdGP63_28_KO and LdGP63_31_KO promastigotes were attached to M□ previously plated on coverslips in a 24-well plate, following the “promastigote attachment assay” protocol. 1:500 dilution of CTB-FITC [Cholera Toxin B subunit (C1655), Sigma] in 1× PBS was added to each well and kept at 4 C for 15 minutes, protected from light. Then the coverslips were washed with 1× PBS, fixed with 2% PFA for 3 min, washed again with 1× PBS, washed with DAPI (Sigma) and mounted with Fluoromount-G (Thermo Scientific) on glass slides. Imaging was done by Leica Stellaris with 63× objective, followed by processing with LAS X software. To estimate the mean fluorescence intensity (CTB) of CTB surrounding each parasite, the ellipse tool of Fiji software was used to first mark the region surrounding each parasite and then “mean” was calculated with the “Measure” tool. To measure the distance between each parasite and its nearest CTB-enriched region, the “Measure” tool was again used and “length” parameter was noted.

### Antibodies

Mouse monoclonal antibodies against Beta-actin (8H10D10), Rab5a (E6N8S), Cas9 (7A9-3A3), mNeonGreen tag (E2S6M) and rabbit monoclonal antibodies against NLRP3 (D4D8T), Caspase-1 (3866S), Cleaved Gasdermin-E (55879S), c-Jun (60A8), mCherry (E5D8F), CD11b (E6E1M) and CD9 (EBL5J) were purchased from Cell Signaling Technology; rabbit monoclonal antibodies against Caspase-3 (A19654), mouse monoclonal antibodies against mCherry tag (AE002) and GST tag (AE001) from Abclonal; rabbit polyclonal antibodies against IL-1β (P420B), Complement C3 (PA521349), mTOR (PA1-518), mouse monoclonal GFP (GF28R) antibody, mouse CD63 monoclonal (MX-49.129.5) antibody and rat monoclonal alpha-tubulin (YL1/2) antibody from Invitrogen; and rabbit polyclonal mCherry antibody (26765) from Proteintech. The secondary HRP-conjugated anti-mouse (A5278) and anti-rabbit (A8275) antibodies were from Sigma Aldrich and SRL. The secondary antibodies for immunofluorescence assays: goat anti-rabbit AlexaFluor 488 (A11008), goat anti-mouse AlexaFluor 488 (A11001), goat anti-rabbit AlexaFluor 594 (A11037), goat anti-mouse AlexaFluor 594 (A11005) were from Invitrogen and goat anti-rat AlexaFluor 647 (44I8) was from Cell Signaling Technology.

### Immunofluorescence assay followed by confocal and structural illumination microscopy

To prepare promastigotes for confocal microscope imaging [76], cultured stationary phase parasites at a number of 1 × 10^4^ were pelleted down in a 1.5 ml centrifuge tube followed by resuspending it in 1× PBS. Then a drop of parasites was layered onto a coverslip and fixed in 2% paraformaldehyde (PFA) for 3 min. Coverslips were then kept in neutralization buffer (0.1 M glycine in PBS). For M□-infected amastigotes, as mentioned in the infection protocol above, coverslips at the experimental time points were collected, rinsed with 1× PBS, fixed with 2% PFA for 3 min and kept in neutralization buffer. The PFA-fixed parasites or M□ were then permeabilized using 0.2% Triton/PBS for 20 min on an orbital shaker. For cell membrane staining, this permeabilization step was omitted. Non-specific binding was blocked with 2% BSA in 0.2% Triton/PBS for another 20 min on an orbital shaker. The parasites or M□ were then incubated with primary antibodies for 1 h, followed by washing and staining with secondary antibody, as required. Further washing with 1× DAPI (Sigma) was done and coverslips were mounted with Fluoromount-G (Thermo Scientific). Imaging was done by Olympus confocal microscopy (FV3000), Leica Stellaris and ZEISS Elyra 7 (Lattice SIM^2^, 60nm) with 63× objective, followed by processing with Fiji software v1.54r.

### Annexin V cell death assay

Parental or KO strains were allowed to infect M□ and incubated for 24 h p.i. with an intermediate 1× PBS wash at 6 h p.i. Initially, the cells were scraped with cell scraper (Nest), pelleted down by centrifuging at 400 *g*, washed with PBS followed by resuspending in FACS buffer (1% NuSera in 1× PBS). However, this protocol resulted in maximum cell death (>80% cell death) in untreated (UI) and parental (P) treated M□ as well, possibly due to mechanical stress of scraping. To minimize the cell death, 10 mM EDTA in 1× PBS (detach solution) was used to detach the cells. Although this recipe did not ensure complete protection of the UI and P infected M□, lower number of stressed near-dead M□ was eventually achieved that differed during every experimental procedure. The cells were submerged in detach solution and incubated at 4°C for 15 min, then the detach solution was vigorously pipetted to dislodge the cells. Detached cells were harvested by centrifuging at 400 *g* at 4°C. Annexin V-FITC (Biolegend) and PI (Thermo Scientific) dyes were added to the cells, diluted as per manufacturer’s protocol, and incubated on ice for 15 min. The presence or absence of Annexin V-FITC or PI was then checked by FACS Aria II system.

### Ultrastructure expansion microscopy

For better visualization, promastigotes were subjected to ultrastructure expansion microscopy [79]. Samples were incubated in an anchoring solution (0.7% formaldehyde and 1% acrylamide in 1× PBS) for 5 h at 37°C. They were then embedded in a 10% acrylamide, 0.1% bis-acrylamide, and 19% sodium acrylate gel, with 0.5% APS and 0.5% TEMED, and allowed to polymerize for 1 h at 37°C. After polymerization, gels were kept in buffer containing 200 mM SDS, 200 mM NaCl, and 50 mM Tris (pH ∼9) at 95°C for 30 min. Gels were subsequently expanded in ultrapure water, immunolabeled as mentioned above, and finally re-expanded in water prior to mounting and imaging.

### Animals

BALB/c mice were maintained and bred under pathogen-free conditions with food supplements and filtered water. All experiments were conducted in accordance with Institutional Animal Ethics Committee (IAEC), IIT Kharagpur guidelines India (IAEC/BI/1 55/2021) and approved by National Regulatory Guidelines issued by Committee for the Purpose of Supervision of Experiments on Animals, Ministry of Environment and Forest, Government of India.

### *In vivo* infection assays

For *in vivo* infections, 1 × 10^7^ lab generated parental-dsRed2 or LdGP63_28_KO-dsRed2 (pXG-dsRed2 plasmid was a kind gift from Dr. Yasuyuki Goto, University of Tokyo) expressing *L. donovani* stationary phase metacyclic promastigotes (suspended in 1× PBS) were injected into healthy female BALB/c mice (4–6 weeks old, 20–25 g each) *via* tail vein route [76]. Each group contained three mice that were sacrificed 12 days and 24 days p.i. Spleens were isolated from the mice and sizes were compared using a standard scale. A small piece of each spleen was stamp-smeared on clean glass slides and later stained with Giemsa solution. Under a sterile hood, the larger parts of the spleens were submerged in NuSera supplemented RPMI 1640 and macerated well on petri plates, followed by RBC lysis with sterile 1× ACK (Ammonium-Chloride-Potassium) lysis buffer and washing with 1× PBS. The macerates were strained with 0.2 µm cell strainer and diluted in FACS buffer. DsRed2+ population was finally measured through flow cytometry.

### Generation of addback strains

For complementing the deleted genes, LdGP63_28 was cloned into pXG-eGFP plasmid (a kind gift from Dr. Yasuyuki Goto, University of Tokyo) in the lab, whereas LdGP63_31 was cloned into pXG-eGFP by Twist Biosciences, both genes upstream of and in the same open reading frame as eGFP. The cloned plasmids were then transfected into LdGP63_28_KO and LdGP63_31_KO promastigotes, with the same protocol as mentioned above, to generate second-copy eGFP-tagged proteases. Post-transfections, parasites were allowed to recover and upon isolating the genomic DNA, the expression of LdGP63_28 and LdGP63_31 was confirmed by PCR with the following primers: LdGP63_28: P35, P36 and LdGP63_31: P37, P38, as mentioned in **Table S4**.

### Synthesis and purification of recombinant LdGP63_28 and LdGP63_31

From the genomic DNA of *L. donovani* promastigotes, previously isolated with DNASure Tissue Mini Kit (Nucleopore), LdGP63_28 (1,701 bp) and LdGP63_31 (1,911 bp) were amplified using the primers (LdGP63_28: P57, P58; LdGP63_31: P59, P60) listed in **Table S4**. Amplified DNAs were run on agarose gels by electrophoresis and target bands were excised from the gels followed by gel purification (Gel and PCR purification Kit, Promega). Cloning of LdGP63_28 or LdGP63_31 into pGEX-6p-1, containing GST tag, was carried out by incubating them with NEBuilder mastermix at 50°C for 1 h. The ligated product was transformed into competent *E. coli* (DH5α) cells followed by plating on ampicillin-LB agar plate, which was incubated overnight at 37°C. Colonies from transformed plate were picked and plasmid was isolated with Surespin Plasmid Miniprep Kit (Nucleopore) to screen for positives by performing PCR with GoTaq polymerase using the above-mentioned primers.

Positive plasmid clones were transformed into *E. coli* Rossetta (DE3) cells by heat shock method and spread on ampicillin-LB agar plates. Colonies from the plate were picked and cultured overnight in LB medium with ampicillin. For secondary culture, 1% from the overnight primary culture was added to it, and allowed to grow at 37°C at 220 rpm continuous shaking. When optical density (OD_600_) values reached 0.6, 1mM IPTG was added to the secondary culture and kept for 5 h with all other parameters intact. The bacteria were then harvested at 3,200 *g* at 4°C, for 15 min, and the recombinant proteins were purified using the previously described protocol [80]. In brief, cell pellet was lysed in PBS containing 1% Triton X-100 and protease inhibitor cocktail followed by freeze-thawing in liquid nitrogen and sonication. After lysis, protein supernatant was collected by spinning sonicated samples at 13,000 *g*. LdGP63_28-GST and LdGP63_31-GST was isolated by using Glutathione-Sepharose beads (Takara Bio), followed by purification by adding reduced glutathione. The purified proteins were separately passed through endotoxin removal column (Pierce, Thermoscientific) and then subjected to Size Exclusion Chromatography (SEC) for further purification. SEC was performed in Superdex 200 Increase 10/300 GL column (AKTA Pure, Cytiva). The column was equilibrated with (50 mM Tris-HCl pH 7.8 and 150 mM NaCl) at a flow rate of 0.5 ml/min, after which samples were loaded and eluted with an additional 1.5 CV. The SEC purified fractions were finally concentrated using Amicon® Ultra-15 Centrifugal Filter Unit (Milipore). The purified proteins were then visualized and confirmed by SDS-PAGE and western blotting (see below), and used for further enzymatic assays.

### SDS-PAGE and western blotting

Samples were prepared by heating in 2× Laemmli buffer for 10□min at 95°C. The samples were subjected to SDS-PAGE system (Bio-Rad) on 8% SDS-polyacrylamide gels. Visualization of proteins on SDS-polyacrylamide gel was done by overnight staining with Coomassie Brilliant Blue R-250 solution (SRL). For western blotting, proteins from the SDS-polyacrylamide gel were transferred to nitrocellulose membrane (GE Healthcare) in the presence of 1× transfer buffer (25 mM Tris, 192 mM glycine, 20% (v/v) methanol) at 20 V for 50 min. The membrane was kept in blocking buffer (5% skim milk powder [SRL] and PBST [1× PBS and 0.05% (v/v) Tween 20]) at 4°C for 1 h, then probed with primary antibody overnight, washed with PBST, incubated with secondary antibody for 2 h, followed by washing with PBST. Bands were visualized in Chemidoc (Biorad) with Chemiluminescent substrate (Cytiva).

### Zymography assay

Gelatine zymography of the purified recombinant proteases was performed using a previously described protocol [31]. Briefly, samples were mixed with loading dye without β-mercaptoethanol and were loaded without boiling on a 10% SDS-PAGE gel containing 1.2 mg/ml of gelatine. Following SDS-PAGE, gels were incubated for 1 h at room temperature in a buffer containing 50 mM Tris pH 7.4, 2.5% Triton X-100, 5 mM CaCl_2_, and 1 µM ZnCl_2_. Gels were then briefly rinsed in distilled water and incubated overnight at 37°C in a buffer containing 50 mM Tris pH 7.4, 5 mM CaCl_2_, and 1 µM ZnCl_2_. Gels were then stained and briefly destained using Coomassie Brilliant Blue (SRL) and visualized in Chemidoc (Biorad).

### Enzyme-substrate assay

The assay was performed by modifying already published protocols [14, 81]. Briefly, M□ were washed thrice with cold PBS containing 1 mM Na_3_VO_4_, scraped off using cell scraper, and lysed on ice by repeated pipetting in 100 μl buffer (50 mM Tris-HCl pH 7.5, 150 mM NaCl, 1 mM EDTA pH 8, 1.5 mM EGTA, 1 mM Na_3_VO_4_, 50 mM NaF, 10 mM Na_4_P_2_O_7_, with protein inhibitor cocktail tablet, Roche). The cells were then collected in a microcentrifuge tube, subjecting the tube to sonication for 10 s burst and 10 s gap for 2 min, centrifuging the tube at 12,000 *g* and collecting the supernatant or the cell lysate. Protein quantification was done with a nanodrop according to the protocol provided by the manufacturer. After protein quantification, the cell lysate was separately added to two different tubes containing LdGP63_28 and LdGP63_31 and incubated at 37°C for 1 h. The tubes were boiled at 95°C for 10 min in Laemmli buffer and migrated in SDS-PAGE gels followed by western blotting.

### Amastigote generation and harvesting

Axenic amastigotes were generated by following a previously published protocol [82], with slight modifications. 1 × 10^4^ promastigotes were harvested and resuspended in M199 medium (pH 5.5) supplemented with 5% FBS. The parasites were allowed to incubate at 22°C with constant shaking for 2 days, and then incubated at 37°C for 2 more days. The parasite culture was then centrifuged at 3,000 *g* and the resultant pellet containing larger promastigotes were discarded. The supernatant was again centrifuged at 7,000 *g* to yield pellet of axenic amastigotes, which were then used for SEM imaging or infecting M□, as required.

For intracellular M□ isolation, M□ plated on 35 mm cell culture dish were allowed to be infected with promastigotes for about 24 h with an intermediate PBS wash for 6 h to remove unbound parasites. At 24 h p.i. the infected M□ were washed with PBS and detached from the dish using cell scraper, followed by repeatedly passing for about 6 times through the needle of a 1 ml syringe. The cells were then centrifuged at 1,200 *g* for 5 min to remove cell debris, and the supernatant collected was centrifuged at 7,000 *g*. The collected amastigote pellet was then used for M□ infection experiments.

### Statistical analysis

Each biological replicate is denoted by n, the exact number for each experiment is mentioned in figure legends. Statistical significance between two groups was determined by two-tailed Student’s *t*-test, while datasets with multiple conditions were analysed by one or two-way ANOVA, followed by Tukey’s or Sidak’s multiple comparisons test (MCT) to assess pairwise differences. A 95% confidence interval (CI) was applied to determine the statistical reliability. Only values of P ≤ 0.05 were considered to be statistically significant, with following notations: ns, non-significant (P > 0.05); *, P ≤ 0.05; **, P ≤ 0.01; ***, P ≤ 0.001; ****, P ≤ 0.0001. All data were represented as mean ± standard error of the mean to illustrate variability within replicates. All statistical analyses were performed with Prism.

## ACKNOWLEDGEMENT

D Manna is funded by Council of Scientific and Industrial Research (CSIR-SRF). S Bhattacharya, S Nandi, D Srinivasan, N Pradhan, and N Pandey are funded by IITKGP *via* GATE fellowship. D Biswas is funded by CSIR-SRF. The work in Yurchenko lab was funded by the EU’s Operational Program ‘Just Transition’ via grant LERCO CZ.10.03.01/00/22_003/0000003.

Authors would like to acknowledge Dr. Tom Beneke (University of Wurzburg, Germany) for his critical inputs and revision during the manuscript preparation. Authors would also like to acknowledge Dr. Supratim Pradhan for his critical inputs throughout the project along with Miss Shubhangi Chakraborty and Miss Aparajita Pati of IDI lab for their help during the experiments; IIR lab, SMST, IITKGP for their help with flow cytometric analysis; confocal microscope facility of Department of Bioscience and Biotechnology (IITKGP), DST-FIST, Govt. of India; confocal facility of SMST (IITKGP) and confocal facility of NISER, Bhubaneshwar, India. Authors would like to thank Miss Puja Kumari and Dr. Dibyendu Samanta, Department of BSBT, IITKGP, for assistance with SEC. Authors would like to thank all the interns of the IDI lab, SMST, especially Miss Pradipti Thakur, Miss Gargi Chatterjee, Miss Anwesha Panja and Miss Sanjana Lahiri, who helped in various experiments related to this project. Computational resources were provided by the e-INFRA CZ project (ID:90254), supported by the Ministry of Education, Youth and Sports of the Czech Republic, and “PARAM Shakti” under the National Supercomputing Mission, IIT Kharagpur, Government of India.

