## Supplementary data for "*Leishmania donovani* GP63 paralogues cooperatively orchestrate visceral infection and persistence"

### SUPPLEMENTARY FIGURES

**Fig S1**

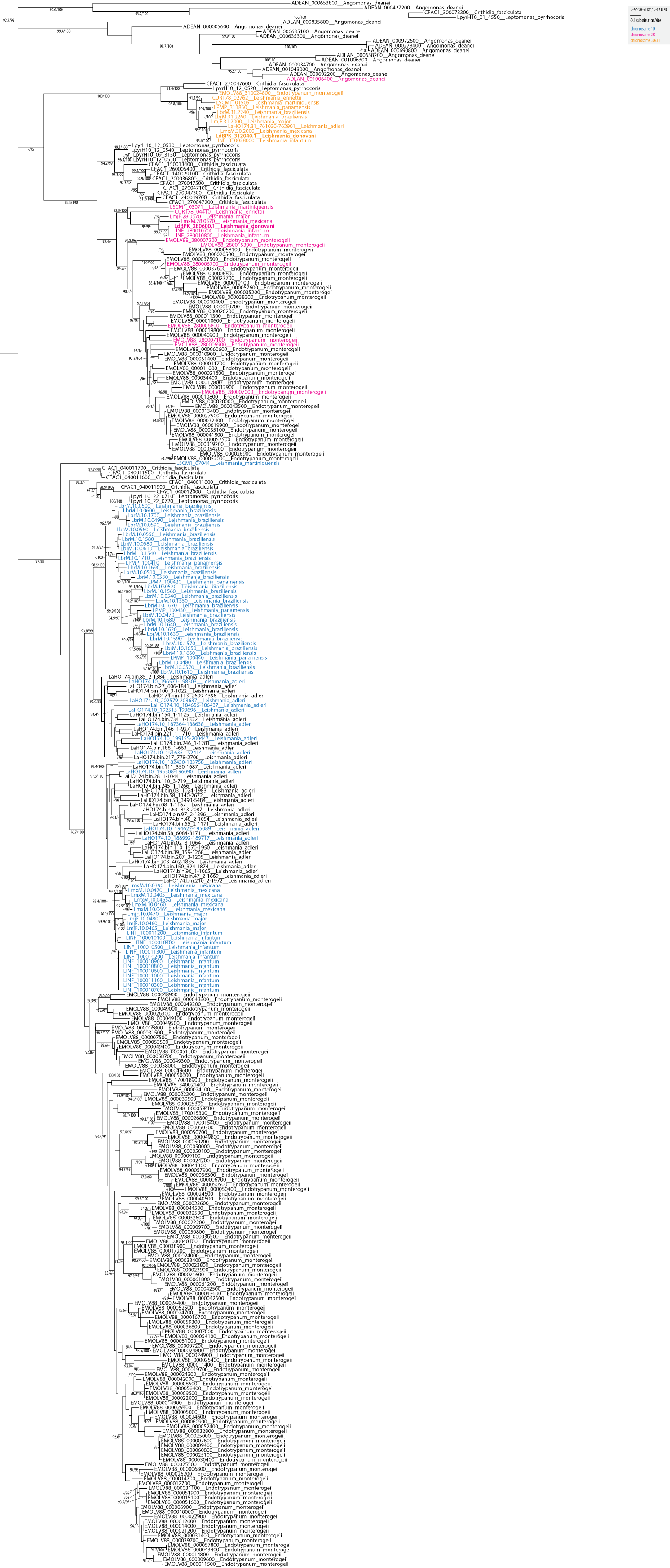

**Figure S1:** **Phylogenetic analysis of selected Leishmaniinae GP63.**

The maximum-likelihood phylogenetic tree was performed in IQ-TREE using PMSF analysis with 1,000 replicates for ultrafast bootstraps (UFB) and Shimodaira-Hasegawa approximate likelihood ratio test (SH-aLRT). Only ≥90 SH-aLRT / ≥95 UFB supports are shown. GP63 proteins encoded on chromosome 10, 28, and 31 are shown in blue, pink, and orange, respectively, as explained in graphical legend.

**Fig S2**
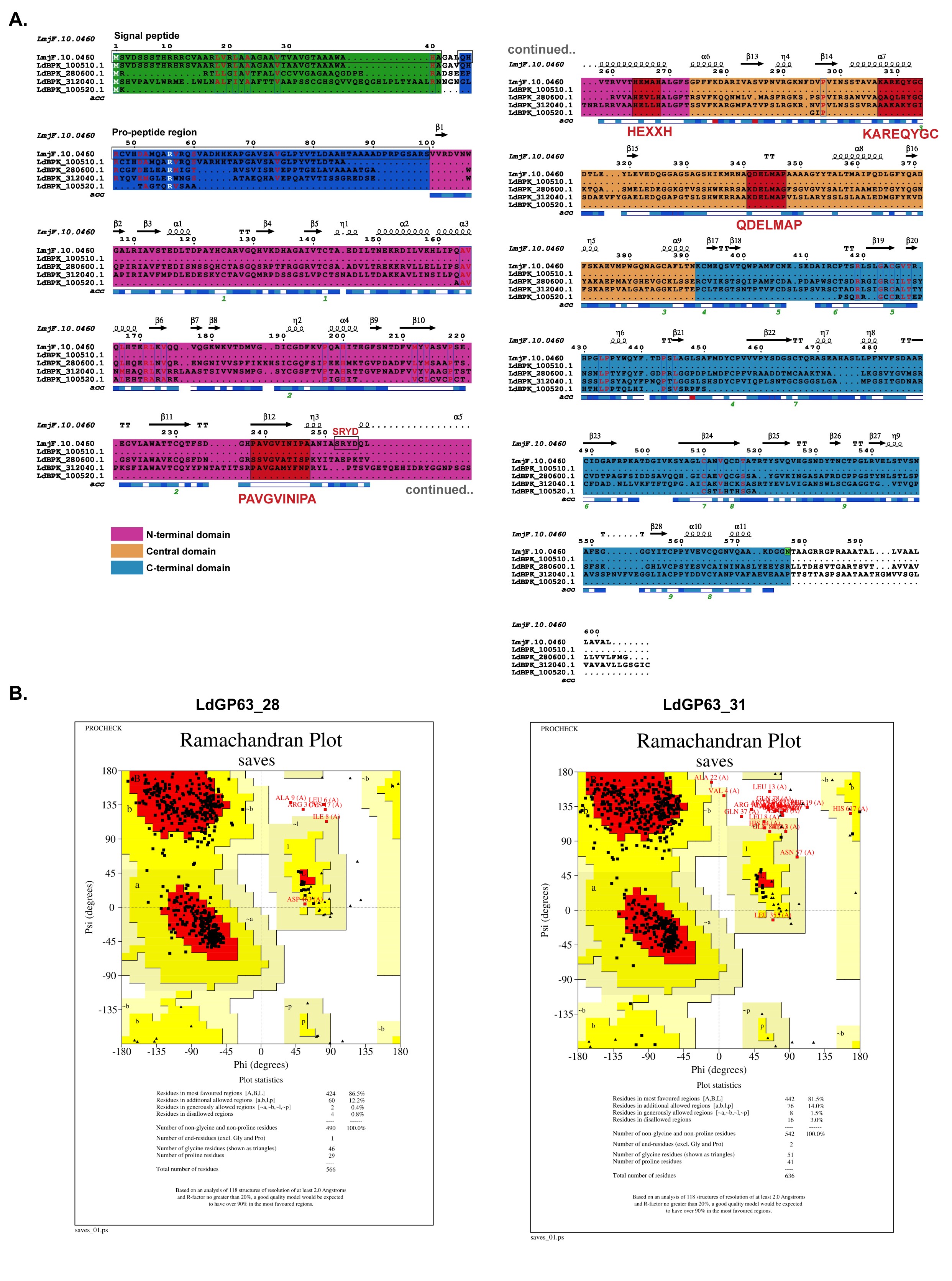

**Figure S2:** **LmGP63, LdGP63_31, and LdGP63_28 show conservation of catalytic and conserved residues.**

A. Multiple sequence alignment of LmGP63 (LmjF.10.0460) and all four LdGP63 sequences- LdGP63_10.51 (LdBPK_100510.1), LdGP63_28 (LdBPK_280600.1), LdGP63_10.52 (LdBPK_100520.1) and LdGP63_31 (LdBPK_312040.1), showing signal peptide (within black box in green region), pro-peptide region (within black box in deep blue region), N-terminal domain (pink), central domain (yellow), and C-terminal domain (blue), with catalytic (HEXXH) and conserved domains (PAVGVINIPA, KAREQYGC, and QDELMAP) boxed in red.

B. Ramachandran plot generated for AlphaFold-modelled LdGP63_28 (left) LdGP63_31 (right) from PROCHECK SAVES server. The distribution of φ and ψ dihedral angles show that most residues lie in favored and allowed regions.

**Fig S3**

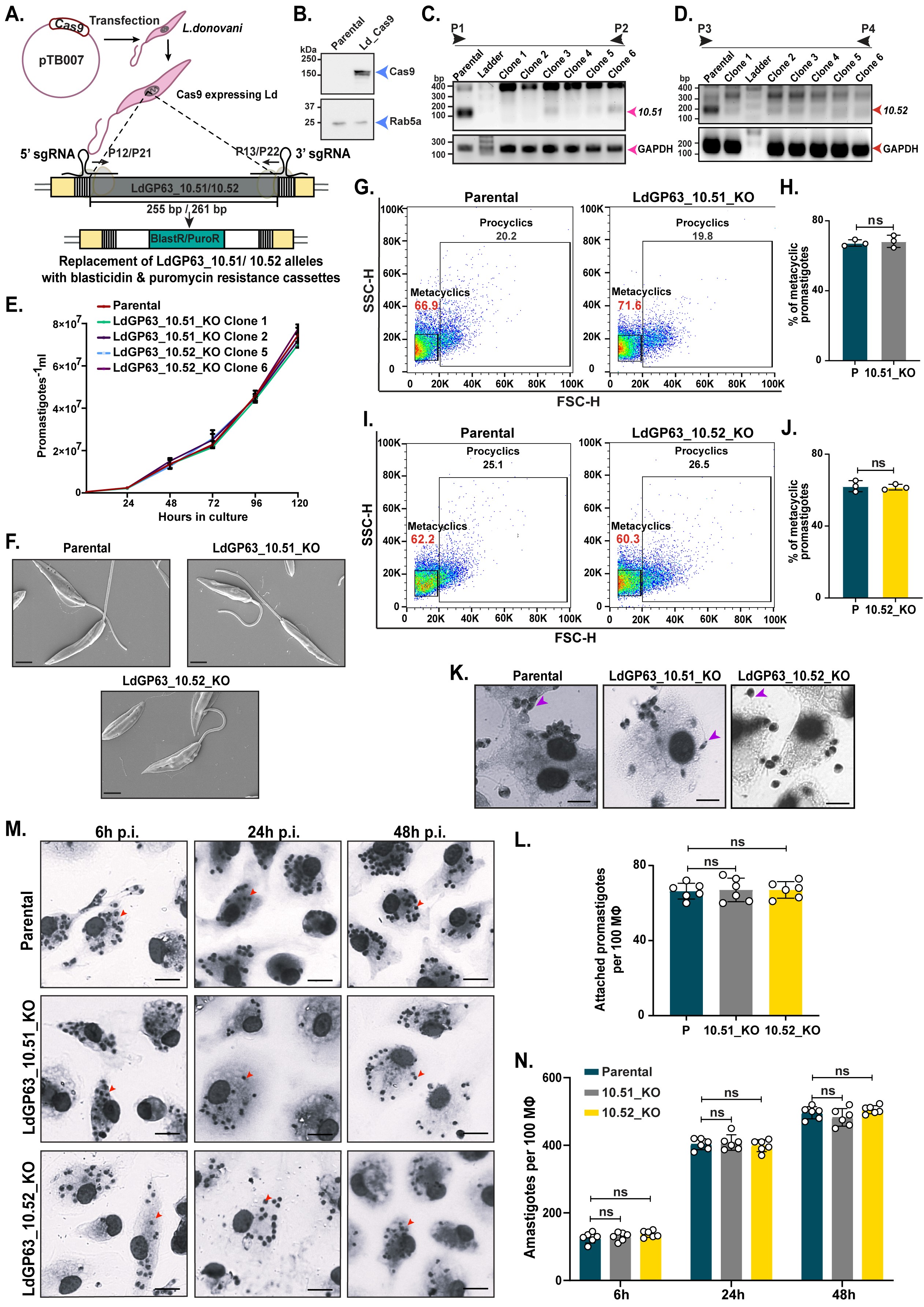

**Figure S3: LdGP63_10.51 and LdGP63_10.52 are not essential proteases for *L. donovani*.**

A. Schematic representation for knock out of LdGP63_10.51 or LdGP63_10.52 with replacement of blasticidin (BlastR) or puromycin (PuroR) resistant cassettes. Primers for BlastR/PuroR cassette insertion in LdGP63_10.51_KO (P12 and P13) and LdGP63_10.52_KO (P21 and P22) are listed in **Table S4**.

B. Western blot confirming Cas9 in pTB007-transfected *L. donovani* lines. Rab5a was used as the loading control.

C-D. PCR confirmation of LdGP63_10.51_KO (C) and LdGP63_10.52_KO (D) from independent clonal populations represented as Clone 1-6. GAPDH was used as the housekeeping control. Primers for LdGP63_10.51_KO confirmation (P1 and P2) and LdGP63_10.52_KO (P3 and P4) are listed in **Table S4**.

E. Growth curve comparing the replicative potential of selected clones (Clone 1 and 2 of LdGP63_10.51, and Clone 5 and 6 of LdGP63_10.52) with respect to parental line.

F. Representative SEM images of log phase parental, LdGP63_10.51, and LdGP63_10.52_KO promastigotes. Scale bar: 2 µm.

G-H. Flow cytometry plot quantifying the number of metacyclics in stationary stage promastigotes of parental and LdGP63_10.51_KO (G) with graphical representation (H) (n = 3).

I-J. Flow cytometry plot quantifying the number of metacyclics in stationary stage promastigotes of parental and LdGP63_10.52_KO (I) with graphical representation (J) (n = 3).

K. Giemsa-stained images comparing initial attachment at 30 min p.i. Attached *L. donovani* promastigotes are pointed by purple arrows. Scale bar: 5 µm.

L. Bar graph showing number of attached parental, LdGP63_10.51_KO, and LdGP63_10.52_KO promastigotes (n = 6).

M. Giemsa-stained images of murine Mϕ infected with parental, LdGP63_10.51_KO, and LdGP63_10.52_KO lines at 6 h, 24 h, and 48 h p.i. Intracellular amastigotes are pointed by red arrows. Scale bar: 10 µm.

N. Bar graph showing the infection and proliferation of parental, LdGP63_10.51_KO, and LdGP63_10.52_KO amastigotes in murine Mϕ at 6 h, 24 h, and 48 h p.i. (n = 6).

Data are presented as mean ± standard error, with each individual point shown. Statistical analysis was performed using paired two-tailed Student’s *t*-test (J), one-way ANOVA followed by Tukey’s MCT (L), and two-way ANOVA followed by Tukey’s MCT (N): ns, non-significant (P > 0.05); *, P ≤ 0.05; **, P ≤ 0.01; ***, P ≤ 0.001; ****, P ≤ 0.0001.

**Fig S4**

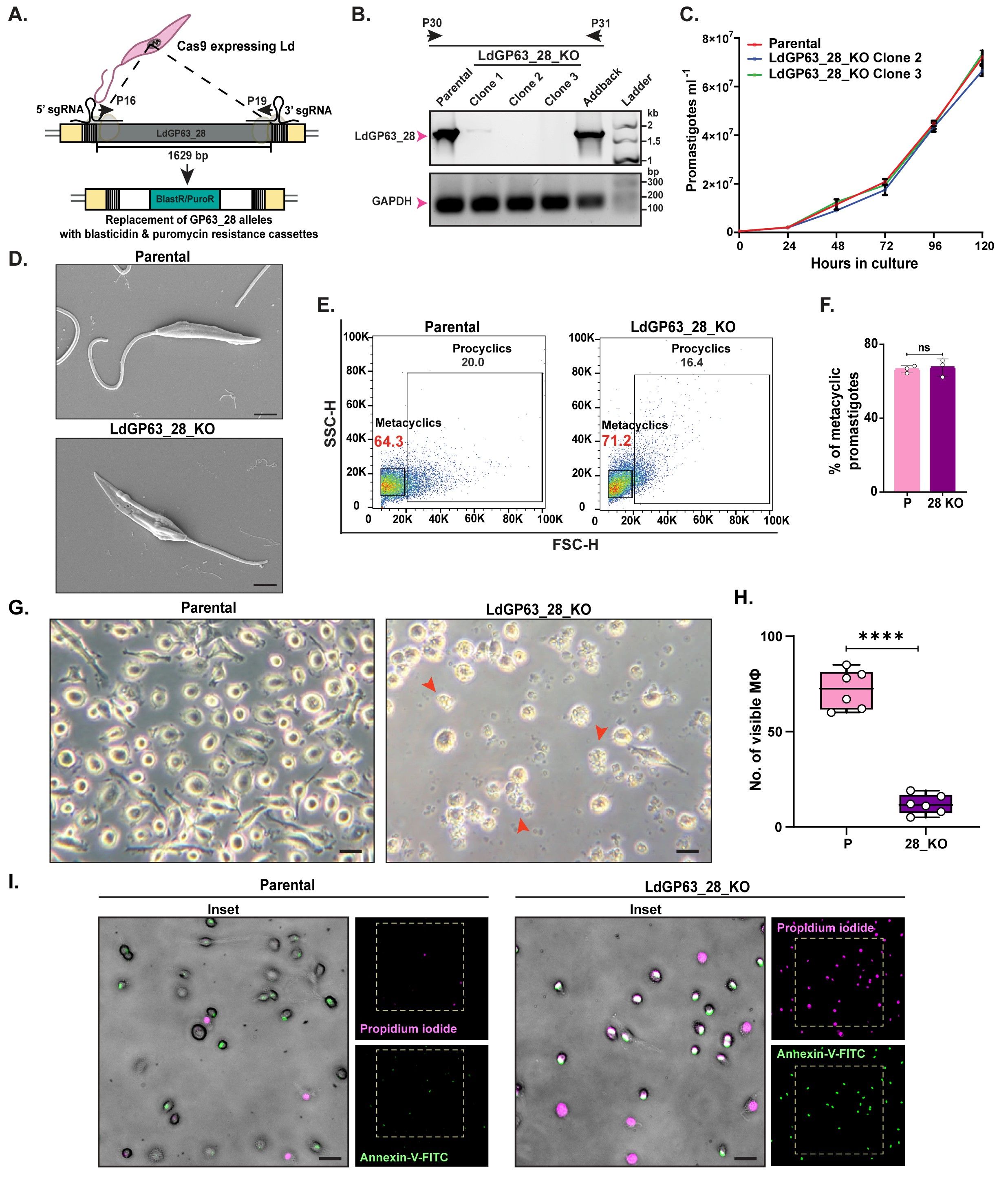

**Figure S4: LdGP63_28 does not affect promastigote survival but affects viability of infected Mϕ.**

A. Schematic representation for knockout generation of LdGP63_28 by replacing both alleles with blasticidin (BlastR) and puromycin (PuroR) resistant cassettes. Primers for BlastR/PuroR cassette insertion in LdGP63_28_KO (P16 and P19) are listed in **Table S4**.

B. PCR confirmation of LdGP63_28_KO in independent clonal populations. Clones 2 and 3 show complete KO of LdGP63_28. Primers for LdGP63_28_KO confirmation (P30 and P31) are listed in **Table S4**.

C. Growth curve comparing the replicative potential of selected LdGP63_28_KO clones (2 and 3) and parental line.

D. Representative SEM images of parental and LdGP63_28_KO promastigotes from log phase culture. Scale bar: 2 µm.

E-F. Flow cytometry plot representing the number of metacyclics in stationary stage promastigotes of parental and LdGP63_28_KO lines (E) with a graphical representation (F).

G-H. Phase-contrast images (G) and their graphical representation (H) depicting murine Mϕ infected with parental and KO parasites at 48 h p.i. (n = 6). Granular, dead cells are indicated by red arrows. Scale bar: 15 µm.

I. Annexin-V-FITC (green) and Propidium iodide (magenta) treated murine Mϕ infected with parental and LdGP63_28_KO parasites at 48 h p.i. Scale bar: 20 µm.

Statistical analysis: ****P ≤ 0.0001; paired two-tailed Student’s t test.

Data are presented as mean ± standard error, with each individual point shown. Statistical analysis was performed using paired two-tailed Student’s *t*-test (F, H): ns, non-significant (P > 0.05); *, P ≤ 0.05; **, P ≤ 0.01; ***, P ≤ 0.001; ****, P ≤ 0.0001.

**Fig S5**

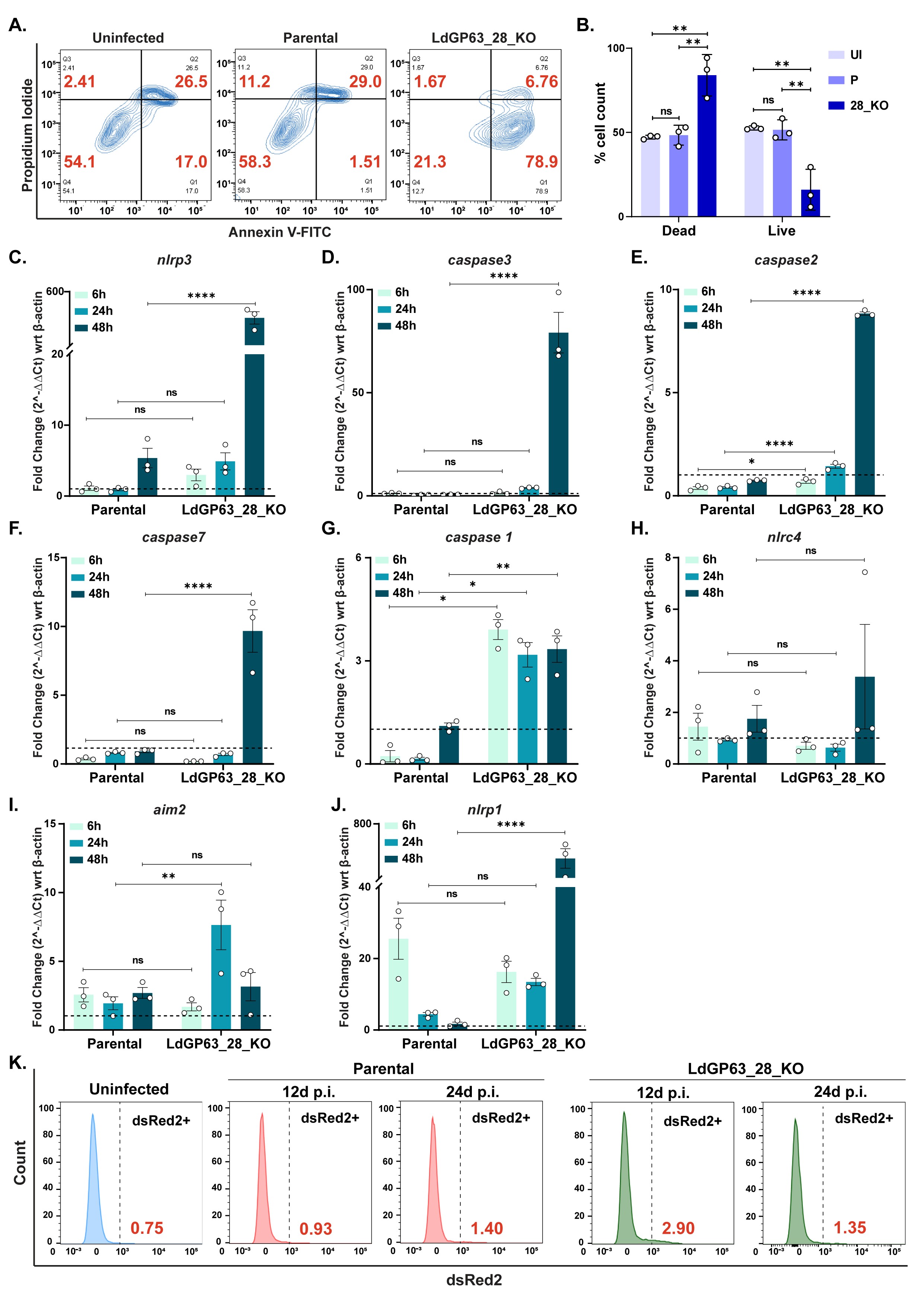

**Figure S5:** **LdGP63_28_KO infected Mϕ show elevated expression of cell death markers.**

A-B. Contour plot showing Propidium iodide vs Annexin V-FITC stained cells comparing uninfected, parental, and LdGP63_28_KO 48 h p.i. Mϕ (A), followed by graph showing the number of dead and live cells after infection (B), (n = 3).

C-J. Relative mRNA expression level of (C) *nlrp3,* (D) *caspase 3,* (E) *caspase 2,* (F) *caspase 7,* (G) *caspase 1,* (H) *nlrc4,* (I) *aim2*, and (J) *nlrp1* of LdGP63_28_KO infected Mϕ at 6 h, 24 h, and 48 h p.i. (n = 3). Dotted horizontal like indicates the uninfected control.

K. Histogram plot of flow cytometric data quantifying the dsRed2+ count in uninfected (blue), parental (red), and LdGP63_28_KO (green) infected mice spleens at 12 days and 24 days p.i. The right side of the vertical dotted line indicates positive dsRed2+ signal with values written in red.

Data are presented as mean ± standard error, with each individual point shown. Statistical analysis was performed using one-way ANOVA followed by Tukey’s MCT (B) and two-way ANOVA followed by Sidak’s MCT (C-J): ns, non-significant (P > 0.05); *, P ≤ 0.05; **, P ≤ 0.01; ***, P ≤ 0.001; ****, P ≤ 0.0001.

**Fig S6**

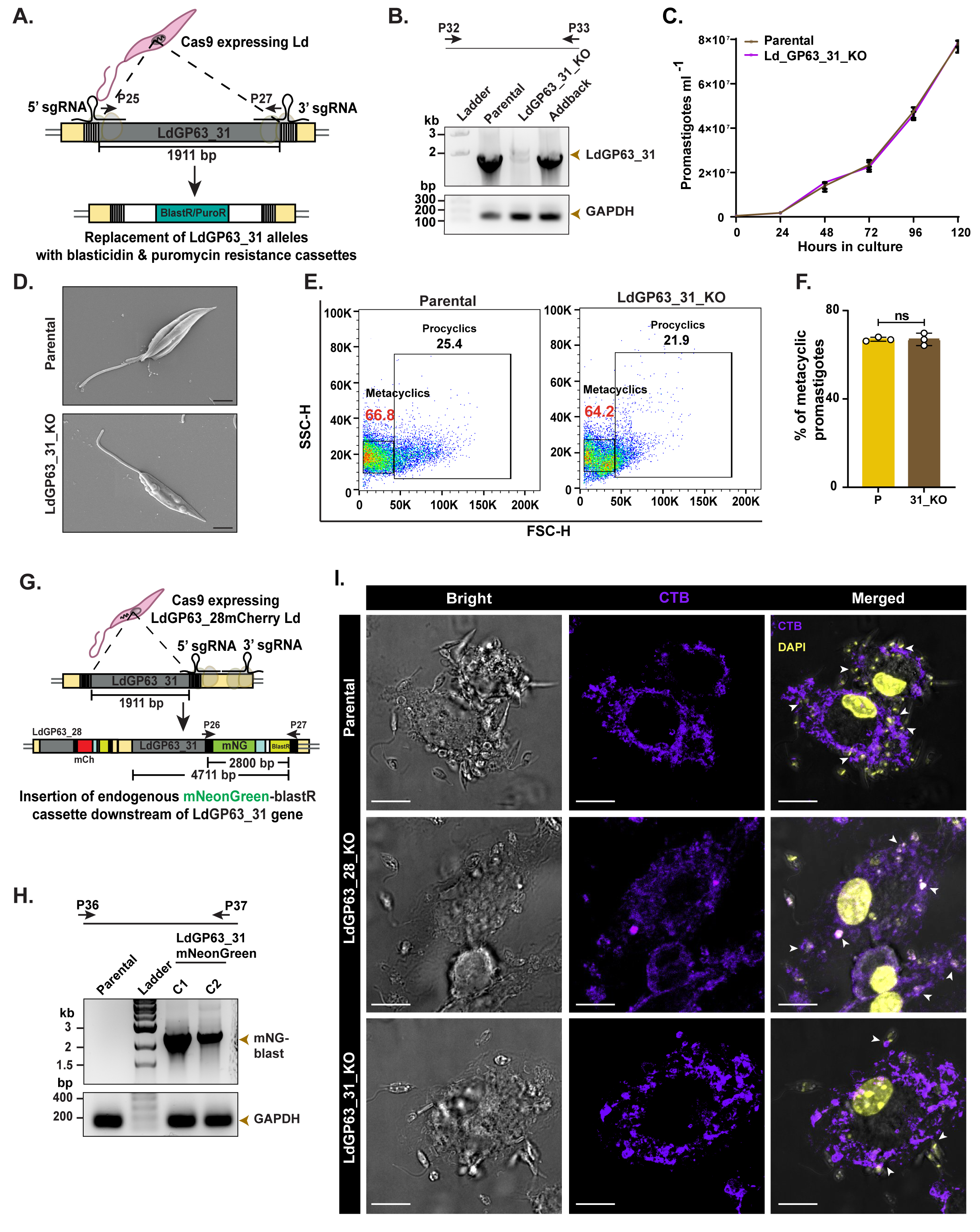

**Figure S6: LdGP63_31 does not affect promastigotes.**

A. Schematic representation for knockout generation of LdGP63_31 by replacing both alleles with blasticidin (BlastR) or puromycin (PuroR) resistant cassettes. Primers for BlastR/PuroR cassette insertion in LdGP63_31_KO (P25 and P27) are listed in **Table S4**.

B. PCR confirmation of KO and addback clones from independent populations. GAPDH was used as the housekeeping control. Primers for LdGP63_31_KO confirmation (P32 and P33) are listed in **Table S4**.

C. Growth curve comparing the replicative potential of selected clones (clone 1 and clone 2) with respect to parental line.

D. Representative SEM images of parental and KO promastigotes in log phase. Scale bar: 2 µm.

E-F. Flow cytometry plot quantifying the number of metacyclics in stationary stage promastigotes of parental and KO (E) with graphical representation (F) (n = 3).

G. Scheme illustrating the endogenous tagging of mNeonGreen-BlastR cassette to the C-terminal of LdGP63_31 in LdGP63_28mCherry-tagged line. Primers for mNGreen cassette insertion downstream of LdGP63_31_KO (P26 and P27) are listed in **Table S4**.

H. PCR confirmation of mNeonGreen cassette insertion using primers (P36 and P37) listed in **Table S4**. GAPDH was used as the housekeeping control.

I. Brightfield (left panel) and confocal micrographs of parental, LdGP63_28_KO and LdGP63_31_KO-infected Mϕ to show the distribution of CTB (middle panel; violet) across the infected Mϕ and its co-localization (right panel) with parasite DAPI (small yellow dots). White arrows indicate the parasite DAPI in close proximity to the host lipid raft. Scale bar: 10 µm.

Data are presented as mean ± standard error, with each individual point shown. Statistical analysis was performed using paired two-tailed Student’s *t*-test (F): ns, non-significant (P > 0.05); *, P ≤ 0.05; **, P ≤ 0.01; ***, P ≤ 0.001; ****, P ≤ 0.0001.

**Fig S7**

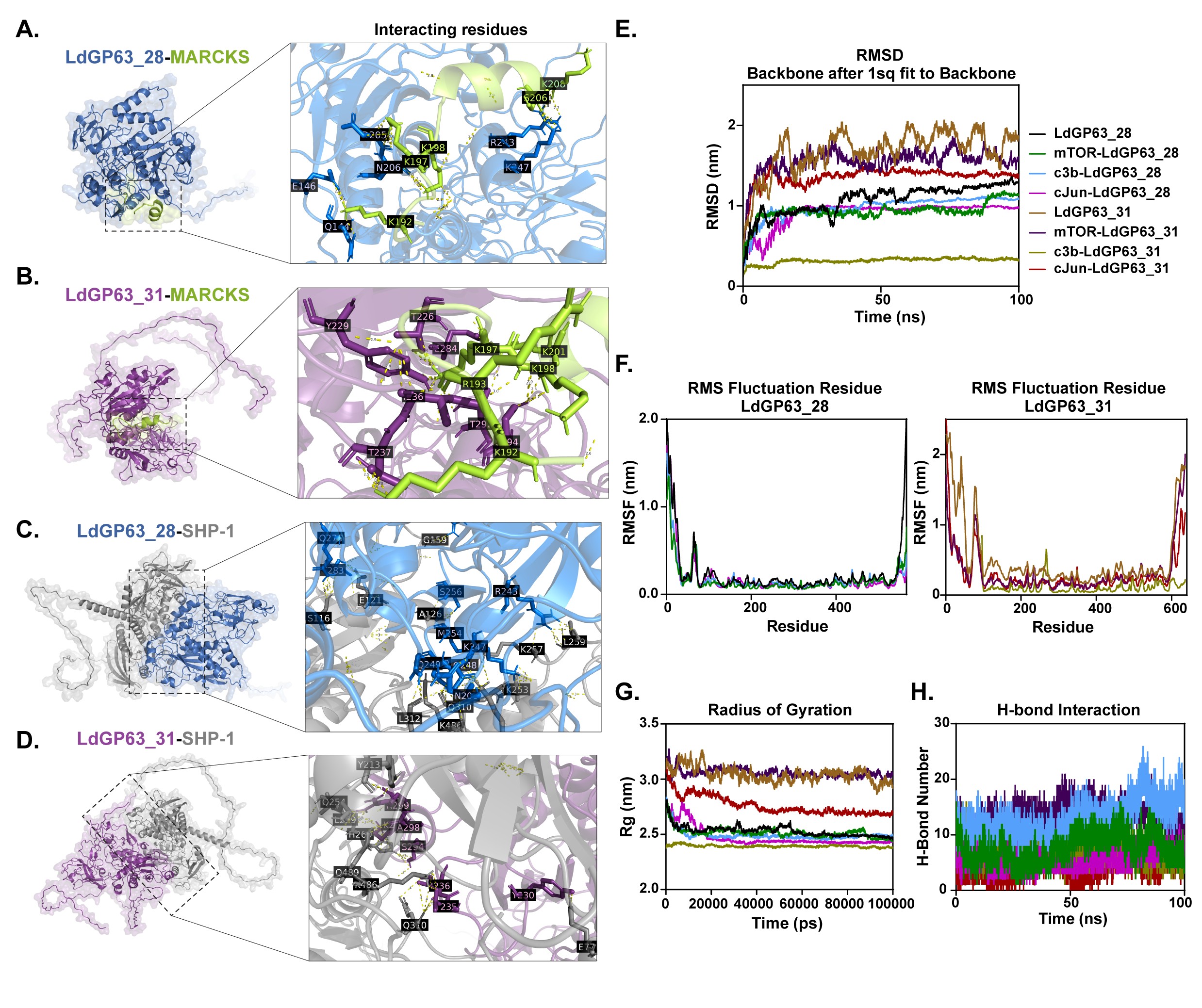

**Figure S7: Docking and molecular dynamics simulation reveals distinct substrate specificities for LdGP63_28 and LdGP63_31.**

A-D. Interactive enzyme-substrate diagrams highlighting the different interacting amino acid residues with their positions in the right-side zoomed structures.

E-H. Graphical plots of (E) root mean square deviation (RMSD in nm), (F) root mean square fluctuation (RMSF in nm), (G) radius of gyration (Rg in nm) and (H) Hydrogen bond numbers obtained from MD simulation of the docked structures. The average ± standard deviation (SD) values are plotted in **Table S3**.

**SUPPLEMENTARY TABLES**

**Table S2: Signal peptide (Sec/SPI) prediction scores of LmGP63 and LdGP63_28**

| **Gene identity** | **Cleavage site position (between amino acid residues)** | **Predicted signal peptide region**  **(cleavage site indicated by “//”)** | **Probability score** |
| --- | --- | --- | --- |
| LmGP63 | Between 41 and 42 | MSVDSSSTHRRRCVAARLVRLAAAGAAVTVAVGTAAAWAHA**//**G | 0.671865 |
| LdGP63_28 | Between 24 and 25 | MRRTLLGIAVTFALVCCVVGAGAA**//**Q | 0.823815 |

**Table S3: Root mean square deviation (RMSD), root mean square fluctuation (RMSF), radius of gyration (Rg), Hydrogen bond numbers (H-bond) scores (average ± standard deviation) obtained from MD simulation of proteases upon interaction with known LmGP63 substrates**

| **Parameter** | **Interaction** | **LdGP63_31** | **LdGP63_28** |
| --- | --- | --- | --- |
| **RMSD (nm)** | Only protease | 1.681 ± 0.2721 | 1.074 ± 0.1865 |
|  | Protease w.r.t. mTOR | 1.552 ± 0.1610 | 0.9324 ± 0.1112 |
|  | Protease w.r.t. cJun | 1.361 ± 0.1247 | 0.9094 ± 0.1605 |
|  | Protease w.r.t. c3b | 0.3174 ± 0.3576 | 0.9804 ± 0.1397 |
| **RMSF (nm)** | Only protease | 0.520404 ± 0.489265 | 0.242896 ± 0.305133 |
|  | Protease w.r.t. mTOR | 0.3286 ± 0.3229 | 0.1670 ± 0.1740 |
|  | Protease w.r.t. cJun | 0.1847 ± 0.2439 | 0.208148 ± 0.291717 |
|  | Protease w.r.t. c3b | 0.1813 ± 0.1912 | 0.35546 ± 0.216012 |
| **Rg (nm)** | Only protease | 3.041 ±0.06558 | 2.530 ± 0.04640 |
|  | Protease w.r.t. mTOR | 3.057 ± 0.03820 | 2.514 ± 0.04369 |
|  | Protease w.r.t. cJun | 2.760 ± 0.08350 | 2.471 ± 0.08840 |
|  | Protease w.r.t. c3b | 2.393 ± 0.01107 | 2.491 ± 0.04458 |
| **H-bond numbers** | Protease w.r.t. mTOR | 13.30 ± 2.278 | 7.531 ± 2.522 |
|  | Protease w.r.t. cJun | 4.013 ± 2.103 | 5.478 ± 1.911 |
|  | Protease w.r.t. c3b | 7.506 ± 2.094 | 13.01 ± 2.476 |
